# The Structural Basis for RNA Binding and Recognition of the Disordered Prion-Like Domain of TDP-43

**DOI:** 10.64898/2025.12.22.695782

**Authors:** Ryan Z. Puterbaugh, Busra Ozguney, Helen L. Danielson, Qizan Chen, Victoria Johnson, Priyesh Mohanty, Jeetain Mittal, Nicolas L. Fawzi

**Affiliations:** Therapeutic Sciences Graduate Program, Brown University, Providence, RI, 02912, USA; Department of Molecular Biology, Cell Biology & Biochemistry, Brown University, Providence, RI 02912, USA; Artie McFerrin Department of Chemical Engineering, Texas A&M College of Engineering, College Station, TX, 77843, USA; Center for Biomedical Engineering, Brown University, Providence, RI 02912, USA; Department of Chemistry, Texas A&M University, College Station, TX, 77843, USA; Interdisciplinary Graduate Program in Genetics and Genomics, Texas A&M University, College Station, TX, 77843, USA

## Abstract

Though the structural details of how RNA interacts with folded RNA-binding domains are well established, how intrinsically disordered regions (IDRs) found in a large fraction of RNA-binding proteins mediate contacts with RNA and if they contribute to binding specificity has not been extensively characterized. The human RNA-binding protein TDP-43 is associated with many RNA processing functions that require its predominantly disordered C-terminal domain (CTD) that forms disease-associated inclusions in ALS, and other neurodegenerative conditions. Here, we demonstrate that TDP-43 CTD directly interacts with RNA primarily via a region of the IDR composed of clustered positively charged residues. Large RNAs act as a multivalent scaffold for CTD monomers, inducing the α-helical segment of TDP-43 CTD to form multimeric protein-protein structures. Additionally, we probe the nucleotide base and amino acid specificity of CTD-RNA interactions, showing that arginine, aromatic and polar residues display a preference for U and G nucleic acid bases over C and A. Finally, we probe the molecular basis for the strong binding interaction between TDP-43 and G4 quadruplex structures and discover similarly avid interactions with cytosine-rich DNA I-motifs. This work deepens our understanding of how disordered regions of proteins contribute to RNA recognition, drive function, and contribute to disease.

**Graphical Abstract:** 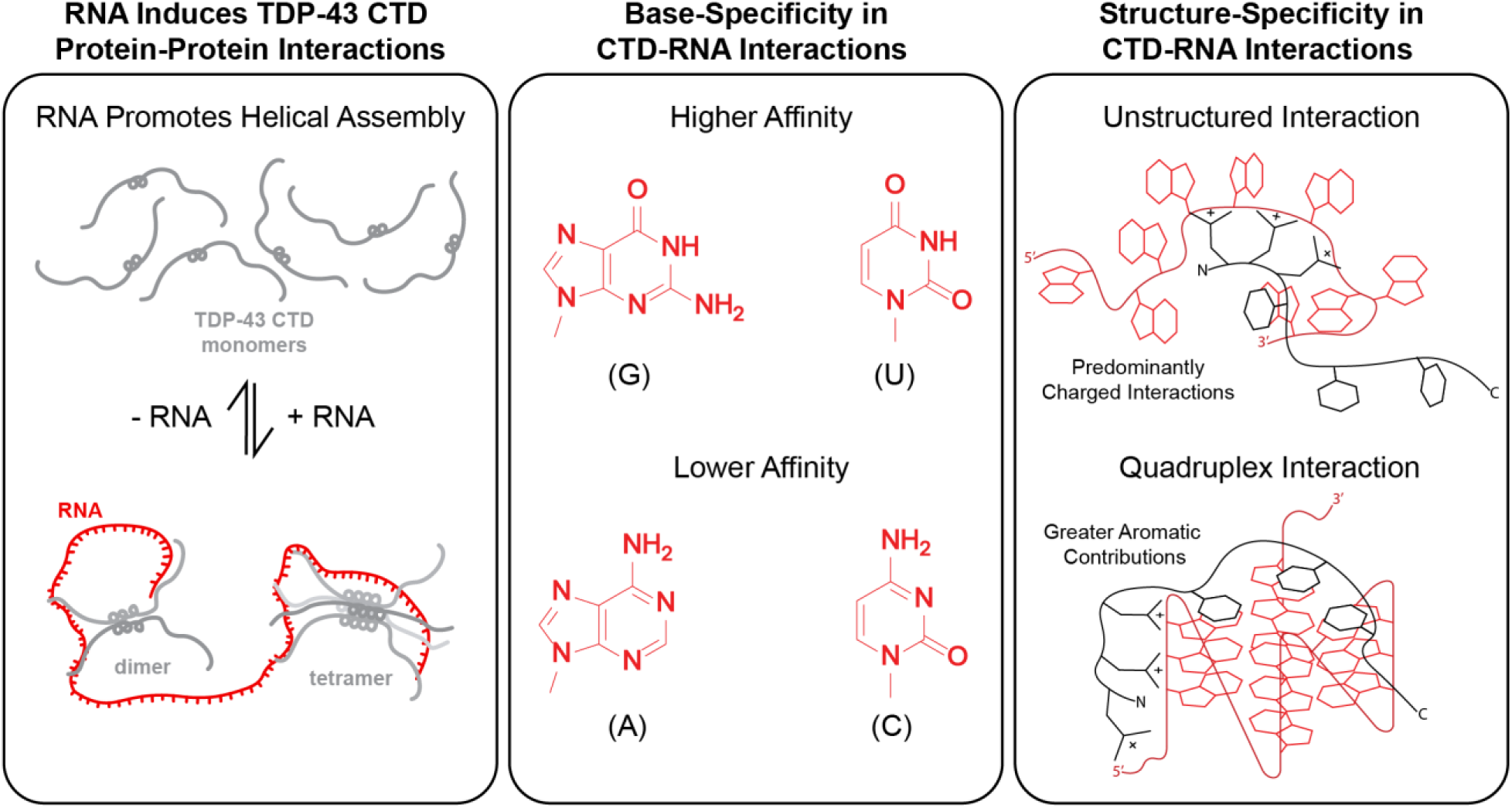

## Introduction

Heterogeneous nuclear ribonucleoproteins (hnRNPs) are a family of RNA binding proteins involved in RNA processing, transport, and localization (1). Structurally, hnRNPs are often composed of multiple canonical RNA-binding domains and long disordered regions that are often aggregation-prone and resemble the sequence composition of yeast prion proteins (i.e. prion-like domains) (1–3). These two structural components are often thought to fulfill distinct roles critical for hnRNP function, with folded RNA-binding domains defining the RNA binding specificity and affinity of hnRNPs (1,3–5), while the disordered regions mediate the ability of hnRNPs to form biomolecular condensates via dynamic protein-protein interactions (6–9). Despite this oversimplified view, numerous examples exist of disordered regions within hnRNPs serving as additional RNA binding regions (10–13), generally occurring through interactions mediated by Arg-Gly-Gly (RGG) or Tyr/Phe-Gly-Gly (Y/FGG) motifs repeated throughout the disordered regions (14–21). Although primarily focusing on the role of these repeats in defining the affinity and/or binding mode to structured RNAs, these studies suggest that disordered regions can make significant contributions to RNA interactions. Interaction scales ranking the energetics of amino acid pair contacts are well studied (22,23), and nucleic acid interaction modes such as base pairing and stacking are well appreciated (24–26). Despite this, investigations into amino acid – nucleic acid interactions in the context of disordered proteins, and if they confer specificity to interactions between disordered region and unstructured RNAs remains understudied (11,25,26). Here, we go beyond to probe the type of dynamic RNA contacts with disordered protein regions expected to be found in biomolecular condensates that may also contribute to hnRNP-RNA binding and recognition.

Trans-activation response DNA-binding protein of 43 kDa (TDP-43) is a ubiquitously expressed hnRNP that plays a role in mRNA transcription, processing, and transport (27–29). Due to its involvement as a causative factor for a number of neurodegenerative diseases such as amyotrophic lateral sclerosis (ALS), frontotemporal dementia (FTD), and the Alzheimer’s disease related dementia known as limbic-predominant age-related TDP-43 encephalopathy (LATE), there has been particular interest in understanding both the normal physiological function and dysregulation of TDP-43 (28,30–32). The structure of TDP-43 consists of a N-terminal domain demonstrated to drive the formation of polymeric assemblies (33), two structured RNA recognition motifs (RRMs) that preferentially bind UG-rich RNA sequences (34–37), and a largely unstructured prion-like C-terminal domain (CTD) which contains a conserved region (CR) adopting transient α-helical structure (38–43). Similar to other hnRNPs (44), this prion-like CTD has been demonstrated to drive TDP-43 phase separation (38), and is aggregation-prone, forming the core of ALS associated inclusions (32,45).

Due to its pathogenic nature, investigations into the structural and cellular mechanisms of TDP-43 CTD phase separation and aggregation have been extensive (38–40,46–52). However, investigations into the role of CTD in TDP-43 interactions with nucleotides and/or RNA have been limited (38,53–58). Unlike other hnRNP disordered regions demonstrated to act as secondary RNA binding domains (14–17), TDP-43 CTD contains only a single true RGG and FGG motif making it unclear if this region in fact interacts with RNA in a consequential manner. We have previously demonstrated that the isolated CTD readily phase separates after addition of RNA (38), implying an interaction with RNA occurs even in the absence of the RRMs. Additionally, other groups have demonstrated via surface plasmon resonance experiments that the CTD alone readily binds to RNA and DNA sequences that form parallel G4 quadruplex structures (55–58). Utilizing a combination of biophysical experiments and atomistic molecular dynamics (MD) simulations in this study, we characterized the structural basis for TDP-43 CTD interactions with RNA and DNA. These results not only broaden our understanding of how TDP-43 recognizes and binds to RNA binding partners, but also how disordered regions of hnRNPs contribute to their role as RNA binding proteins.

## Methods

### Constructs

The following constructs were transformed into and cultured in BL21 Star (DE3) *Escherichia coli* (Life Technologies) for protein expression:

- TDP-43 CTD (Addgene ID: 98669)
- TDP-43 CTD A326P (Addgene ID: 98672)
- TDP-43 CTD M337P (Addgene ID: 98675)
- TDP-43 CTD 3RtoK (R268K-R272K-R275K) (Addgene ID: 250594)
- TDP-43 CTD 3RtoA (R268A-R272A-R275A) (Addgene ID: 250589)
- TDP-43 CTD allR/KtoA (R268A-R272A-R275A-R293A-R361A-K408A) (Addgene ID: 250607)
- TDP-43 CTD 2R/KtoA (R293A-R361A-K408A) (Addgene ID: 250608)

### Protein Expression and Purification

TDP-43 CTD constructs were expressed in either LB or M9 minimal media supplemented with ^15^NH_4_Cl following a slightly modified version of a previously described protocol (38). In brief, 1 L bacterial cultures were grown to a OD_600_ between 0.7-0.8 before being induced with 1 mM IPTG for 4 hours at 37°C. Cells pellets were harvested by centrifugation (6000 rpm, 15 min, 4°C) and stored at −80°C. Cell pellets were resuspended in 25 mL of buffer (20 mM Tris, 500 mM NaCl, 10 mM imidazole, pH 8.0) followed by lysis using a ultrasonic cell disruptor. Inclusion bodies were pelleted via centrifugation (20,000 rpm, 1 hour, 4°C) resuspended in 23 mL of solubilizing buffer (20 mM Tris, 500 mM NaCl, 10 mM imidazole, 8 M urea, pH 8.0) then cleared of cellular debris via a second centrifugation (25,000 rpm, 1 hour, 4°C). The resulting supernatant was filtered and loaded onto a 5 mL Histrap HP column after which bound protein was eluted using a 10-500 mM linear imidazole gradient over 5 column volumes.

Fractions containing TDP-43 CTD were then pooled and desalted using a HiPrep 26/60 Desalting Column into a TEV cleavage buffer (20 mM Tris, 500 mM GdnHCl, pH 8) and subjected to an overnight TEV cleavage at room temperature. Following TEV cleavage solid urea was added to achieve an 8 M concentration before the protein being once again loaded onto a 5 mL Histrap HP column to remove TEV and now cleaved 6xHis tag. Fractions containing cleaved TDP-43 CTD were then pooled, exchanged into a storage buffer (20 mM MES, 8 M Urea, pH 6.1), and concentrated to approximately 1.5-2 mM at which point 100 uL aliquots were flash frozen and stored at −80°C.

### RNA and DNA

Ribonucleic acid from torula yeast (Product # R6625), polyU potassium salt (Product # P9528), polyC potassium salt (Product # P4903), polyA potassium salt (Product # P9403), and polyG potassium salt (Product # P4404) were all purchased as lyophilized powder from Millipore Sigma. Before use, all RNAs were resuspended in either 20 mM MES buffer pH 6.1 or 20 mM sodium phosphate buffer pH 6.75 then desalted into the same buffer using a 2 mL Zeba spin desalting column (Thermo Scientific, Product # 89890) according to manufacturer instructions after which they were aliquoted and frozen at −20°C. ssDNA oligos were purchased from Genscript as desalted dry powder with no 5’ or 3’ modifications. ssDNA oligos were dissolved in 20 mM MES buffer pH 6.1 to a stock concentration of 1 mM before use. The sequences of oligos are as follows:

- dT_12_ TTTTTTTTTTTT
- dU_12_ UUUUUUUUUUUU
- dA_12_ AAAAAAAAAAAA
- dG_12_ GGGGGGGGGGGG
- dC_12_ CCCCCCCCCCCC
- c-Myc G4 GCTTATGGGGAGGGTGGGGAGGGTGGGGAAGGTG
- c-Myc I-motif CACCTTCCCCACCCTCCCCACCCTCCCCATAAGC
- (TG)_17_ TGTGTGTGTGTGTGTGTGTGTGTGTGTGTGTGTG
- (AC)_17_ ACACACACACACACACACACACACACACACACAC

### Microscopy

Microscopy samples were prepared by dilution of TDP-43 CTD stocks to a final concentration of 20 μM in either a 20 mM MES 160 mM urea pH 6.1 buffer or 20 mM sodium phosphate 160 mM urea pH 6.75 buffer supplemented with 0 mg/mL, 0.05 mg/mL, 0.5 mg/mL, and 3 mg/mL yeast RNA. Samples were thoroughly mixed before spotting 10 μL onto glass coverslips for imaging. DIC micrographs were obtained for each sample using a Nikon Ti2-E Microscope equipped with a 60x oil objective and 1.5x digital enhancement. The resulting images were processed using Fiji software.

### Turbidity Measurements

TDP-43 CTD variants were diluted to a final concentration of 20 μM in a 150 μL volume of in either a 20 mM MES 160 mM urea pH 6.1 buffer or 20 mM sodium phosphate 160 mM urea pH 6.75 buffer supplemented with 0 mg/mL, 0.015 mg/mL, 0.03 mg/mL, 0.045 mg/mL, 0.06 mg/mL, 0.09 mg/mL, 0.15 mg/mL, 0.3 mg/mL, 0.45 mg/mL, 0.9 mg/mL, and 3 mg/mL RNA. Samples were thoroughly mixed following dilution before 50 μL aliquots were loaded onto a Greiner bio-one 96 well plate. The 600 nm absorbance of wells were then immediately measured using a Cytation 5 plate reader.

### NMR Spectroscopy

#### Solution NMR Samples and RNA Titrations

All NMR experiments were performed using Bruker Avance 850 MHz or 600 MHz ^1^H Larmor frequency spectrometers with HCN TCI z-gradient cryoprobes at 298K. NMR-titrations of TDP-43 CTD constructs with RNA were performed by desalting ^15^N labeled protein samples into in either a 20 mM MES pH 6.1 buffer or 20 mM sodium phosphate pH 6.75 buffer using 0.5 mL Zeba spin desalting columns (Thermo Scientific, Product #89882), before being diluted to a final concentration of 20 μM in buffer supplemented with 5% D_2_O and the appropriate concentration of RNA for each titration point. ^1^H-^15^N HSQCs were collected of samples with 200* and 2048* complex pairs in the indirect ^15^N and direct ^1^H dimensions with acquisition times of 105 ms and 200 ms and sweep widths of 22.0 ppm and 12.0 ppm centered around 116.5 ppm and 4.7 ppm, respectively. NMR spectra were processed with NMRPipe (59) and analyzed with NMRFAM-Sparky (60). Resonance assignments for WT TDP-43 CTD, TDP-43 CTD A326P, and TDP-43 CTD M337P were transferred from the Biological Magnetic Resonance Bank (BMRB) records (BMRB 26823, BMRB 26828, and BMRB 26831). Resonance assignment of mutant variants were performed primarily by overlay with the WT. Residues near mutation sites that had undergone dramatic chemical shifts were assigned by estimation of their new position using the Poulsen IDP/IUP random coil chemical shifts calculator (https://spin.niddk.nih.gov/bax/nmrserver/Poulsen_rc_CS/) (61,62). Peak Intensity ratios were calculated by normalizing the peak height of residues in the presence of RNA to the peak height in the absence of RNA. The ^1^H and ^15^N chemical shifts (Δδ^1^H and Δδ^15^N) were calculated as the difference in the chemical shift in the presence of RNA minus the chemical shift in the absence of RNA. The combined chemical shift (Δδ^1^H-^15^N) was calculated using equation 1.

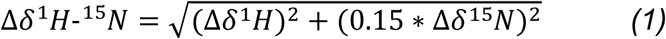

#### NMR Transverse Relaxation

^15^N *R*_2_ of WT CTD in the absence and presence of 3 mg/mL yeast RNA was measured at 850 MHz ^1^H using a standard pulse sequence (hsqct2etf3gpsitc3d from Bruker Topspin 3.2) acquired with 128* and 1536* complex pairs in the indirect ^15^N and direct ^1^H dimensions with acquisition times of 78 ms and 150 ms and sweep widths of 19.0 ppm and 12.0 ppm centered around 116.5 ppm and 4.7 ppm, respectively. ^15^N *R*_2_ experiments had an interscan delay of 2.5 s, a Carr-Purcell-Meiboom-Gill (CPMG) field of 556 Hz, and total *R*_2_ relaxation CMPG loop-lengths of 16.3 ms, 32.6 ms, 48.9 ms, 65.2 ms, 163.0 ms, 195.6 ms, and 244.5 ms. ^1^H_N_ *R*_2_ of WT CTD in the absence and presence of 3 mg/mL yeast RNA was measured at 850 MHz ^1^H with 128* and 1536* complex pairs in the indirect ^15^N and direct ^1^H dimensions with acquisition times of 78 ms and 172 ms and sweep widths of 19.0 ppm and 12.0 ppm centered around 116.5 ppm and 4.7 ppm, respectively. Each ^1^H_N_ *R*_2_ experiment comprised six interleaved ^1^H_N_ *R*_2_ relaxation delays: 0.2 ms, 10.2 ms, 20.2 ms, 40.2 ms, 60.2 ms, 90.2 ms.

#### Diffusion-Order NMR Spectroscopy

Protein diffusion in the presence and absence of ssDNA was measured using a standard diffusion-weighted one-dimensional ^1^H pulse sequence (ledbpgppr2s from Bruker Topspin 3.2) with 16 gradient amplitudes linearly spaced between 2% and 98% of the maximum gradient strength (49 G/cm). Each spectrum was acquired with 3072 points, an acquisition time of 172 ms, and a sweep width of 10.5 ppm centered on 4.7 ppm. For all experiments the diffusion delay (Δ) was set to 0.1 s and the net diffusion encoding pulse width (*δ*) was set to 3.6 ms. The apparent diffusion time was calculated by integrating the peak area of amino acid side chain protons from 1.945-1.995 ppm using Bruker Topspin 4.4. The peak area at each gradient strength (*I_G_*) was then plotted as a function of gradient strength (*G*) and fitted to equation 2 using the curve_fit function from the scipy.optimize library (version 1.10.0).

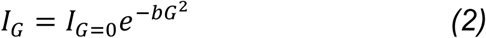

The parameter *b* from equation 2 is defined by the diffusion coefficient (*D*), the gyromagnetic ratio (*γ*), diffusion delay (Δ), and net diffusion encoding pulse width (*δ*) as described in equation 3.

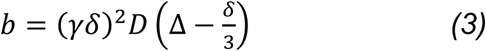

Given that the gyromagnetic ratio, diffusion delay, and net diffusion encoding pulse width are all constant between titration points, the only parameter that has an effect on the value of *b* between titration points is the change in the apparent diffusion coefficient. To better represent how the diffusion coefficient changed as a result of ssDNA addition, we decided to calculate a normalized diffusion coefficient (*D_norm_*) for each titration point in reference to the diffusion coefficient of the protein in the absence of ssDNA using equation 4.

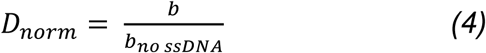

#### Circular Dichroism Spectroscopy of RNA Constructs

12mer ssDNA oligos were diluted from 1 mM stocks to final concentrations of 60 μM in 20 mM MES buffer pH 6.1, while c-Myc G4 and c-Myc I-motif oligos where diluted to 20 μM in 20 mM MES buffer pH 6.1. CD spectra were collected using a JASCO J-815 spectropolarimeter using a 2 mm path length quartz cell at 20°C. Each sample was scanned three times to obtain an average spectrum from 190 nm to 350 nm.

### Molecular Dynamics (MD) Simulation

#### System Preparation

System preparation for all-atom simulations was conducted in GROMACS (63). For modelling the protein, Amber03ws force field paired with TIP4P/2005 water model was chosen due to its optimized protein-water interactions that prevent overly compact states in intrinsically disordered regions (64,65). RNA molecules were parameterized using the χOL3 force field with refined glycosidic torsion angles (66) and adjusted phosphate oxygen radii (67). The compatibility of the force fields used in this study has been previously demonstrated (37).

Initial configurations of protein were obtained from previously generated all-atom simulations performed in our group (unpublished data) and AlphaFold prediction (AF-Q13148-F1-v4). For the RNA component, 12-nucleotide unstructured RNA strands with varying sequences were generated using the Nucleic Acid Builder (NAB) module implemented in AmberTools (68,69). 10 RNA oligo molecules were randomly placed around the protein using the “gmx insert” utility in GROMACS. This approach increases the probability of protein-RNA encounters without imposing artificial constraints on the binding mode. Each system was subsequently solvated in a truncated octahedral water box with a 12.5 nm box length. Counter ions were added to maintain electroneutrality at a physiological salt concentration of 0.10 M NaCl, employing the improved ion parameters developed by Luo and Roux (70). The GROMACS topology and coordinate files were then converted to AMBER format (parm7 and rst7) using the “gromber” utility in ParMED (71). During this conversion, hydrogen mass repartitioning (72) to 1.5 amu was implemented, enabling an extended 4 fs timestep in production simulations.

#### Simulation Protocol

All production simulations were performed in Amber22 (68) at 1 bar and 300 K following a multi-step equilibration process. First, a minimization run was carried out using the steepest descent algorithm for 5000 cycles followed by the conjugate gradient algorithm for an additional 2500 cycles, with all non-hydrogen atoms of the protein restrained by a 5 kcal/(mol*Å^2^) force constant. Next, the system was subjected to a 2 ns heating protocol with the restraint force constant reduced to 0.5 kcal/(mol*Å^2^), where temperature increased linearly from 0 K to 300 K during the first 0.5 ns and was maintained at 300 K thereafter. An additional 10 ns simulation under canonical ensemble (NVT) was conducted without any restraints, followed by 10 ns under isothermal-isobaric ensemble (NPT) for pressure equilibration. Temperature control was achieved using Langevin dynamics with a 1.0 ps^-1^ collision frequency, while pressure was regulated using a Monte Carlo barostat (73) with isotropic coupling, 1.0 ps relaxation time, and volume change attempts occurring every 100 steps. SHAKE algorithm (74) was employed to constrain bonds involving hydrogen atoms. A non-bonded cutoff of 0.9 nm was applied for short-range interactions, and long-range electrostatic interactions were treated using the Particle Mesh Ewald (PME) (75) method. Finally, each system without RNA and with varying RNA sequences was subjected to three 4 µs NPT production runs using identical parameters as the final step of the equilibration phase.

To decide the equilibration time and confirm that the overall length of our individual trajectories (4 μs each) allows the conformational exploration beyond the initial configurations, we calculated radius of gyration (R_g_) (Figure S4A and S7), a well-established measure used for evaluating chain global dimensions. By calculating the time autocorrelation function of R_g_ over each of these trajectories, we analyzed the timescale of chain relaxation. Based on this analysis, 500 ns was selected as equilibration time for all systems.

#### All-Atom Dense Phase Molecular Dynamics (MD) Simulation

Initially, TDP-43 CTD and RNA chains (AUG12 sequence, 12-nucleotide long) were placed in a box of 10.5 nm x 10.5 nm x 73.5 nm and simulated using HPS Urry model (76) for proteins and HPS model (77) for RNA. Residues 320–340 of the TDP-43 CTD were constrained to maintain a rigid α-helical conformation, as our previous study identified this helix as a key determinant of TDP-43 CTD phase separation at 300 K. Following equilibration, the proteins formed a condensed phase, whereas several RNA molecules remained in the dilute phase. To improve computational efficiency in all-atom simulations, we trimmed the system by keeping only RNA molecules that entered the TDP-43 CTD condensate. This yielded a backmapped all-atom system comprising 40 CTD and 11 RNA chains to all-atom configurations using MODELLER (78) and Arena (79) software respectively. The atomistic slab was then prepared for simulations in GROMACS. The parameters used for protein and RNA were identical to those used for dilute phase simulations. System was solvated using TIP4P/2005 water molecules in 10.5 nm x 10.5 nm x 40.0 nm simulation box, and 0.10 M NaCl which was modeled with Luo and Roux (70) parameters, was added. Energy minimization was first performed in GROMACS until the maximum force on any atom was below 1000 kJ/mol/nm. The system was then equilibrated by running 100 ps of NVT followed by 100 ps of NPT. During equilibration, the temperature was maintained at 300 K using the velocity-rescale (V-rescale) thermostat (80), and the pressure was controlled at 1 bar using the Parrinello–Rahman barostat (81,82) to equilibrate the system density. Production runs were performed in Amber22 after converting GROMACS input files to AMBER format. Monte Carlo barostat was used for pressure coupling. Hydrogen involving bonds were restrained with SHAKE algorithm, while long-range electrostatic interactions were computed using the PME method and van der Walls interactions were truncated at 0.9 nm. 1 µs NPT production runs at 1 bar and 300 K with and without RNA were conducted.

#### Structural and Contact Analysis

All contact analysis was conducted using the MDAnalysis “neighbor search library” and/or “distance” module (83,84). A van der Waals interaction was considered formed when any heavy atom of the protein was within 4.5 Å of any heavy atom of RNA molecule, and these interactions were added to the pairwise contact probability. The contact propensity of each protein residue with RNA (1D contact map) was calculated by summing all contact instances across the trajectory and then normalizing by the number of RNA molecules (n=10) present in the simulation. For residue type contact maps, contact probabilities of amino acid type pairs were summed across all instances. GROMACS (63) utility tools were used for calculations of R_g_ and radial distribution function (RDF).

## Results

### TDP-43 CTD - RNA Interactions are Primarily Mediated by N-Terminal Arginine Residues

Our previous attempts to characterize TDP-43 CTD interactions with RNA by solution NMR failed due to RNA inducing phase separation of the CTD resulting in poor signal to noise as the majority of the protein localizes to droplets consisting of a condensed protein-RNA phase (i.e. a biphasic sample) that results in slowed motions and challenging spectral properties (85). RNA-mediated reentrant phase separation, a biophysical phenomenon in which RNA strongly induces phase separation of proteins at low concentrations but begins to dissolve droplets at high concentrations, has been described for phase separating proteins and peptides (86,87). Inspired by a similar strategy using peptides to prevent phase separation for NMR studies (88), we hypothesized that TDP-43 CTD may also undergo reentrant phase separation at higher RNA concentrations than previously tested, thereby allowing characterization of CTD-RNA interactions in the “NMR visible” dispersed phase. We observed via differential interference contrast (DIC) microscopy (**Fig. 1A**) and optical turbidity (**Fig. 1B**) that low concentrations of torula yeast RNA strongly induced phase separation, but higher RNA concentrations dissolved these droplets, indicating that the protein undergoes RNA-mediated re-entrant phase separation.

**Figure 1:**
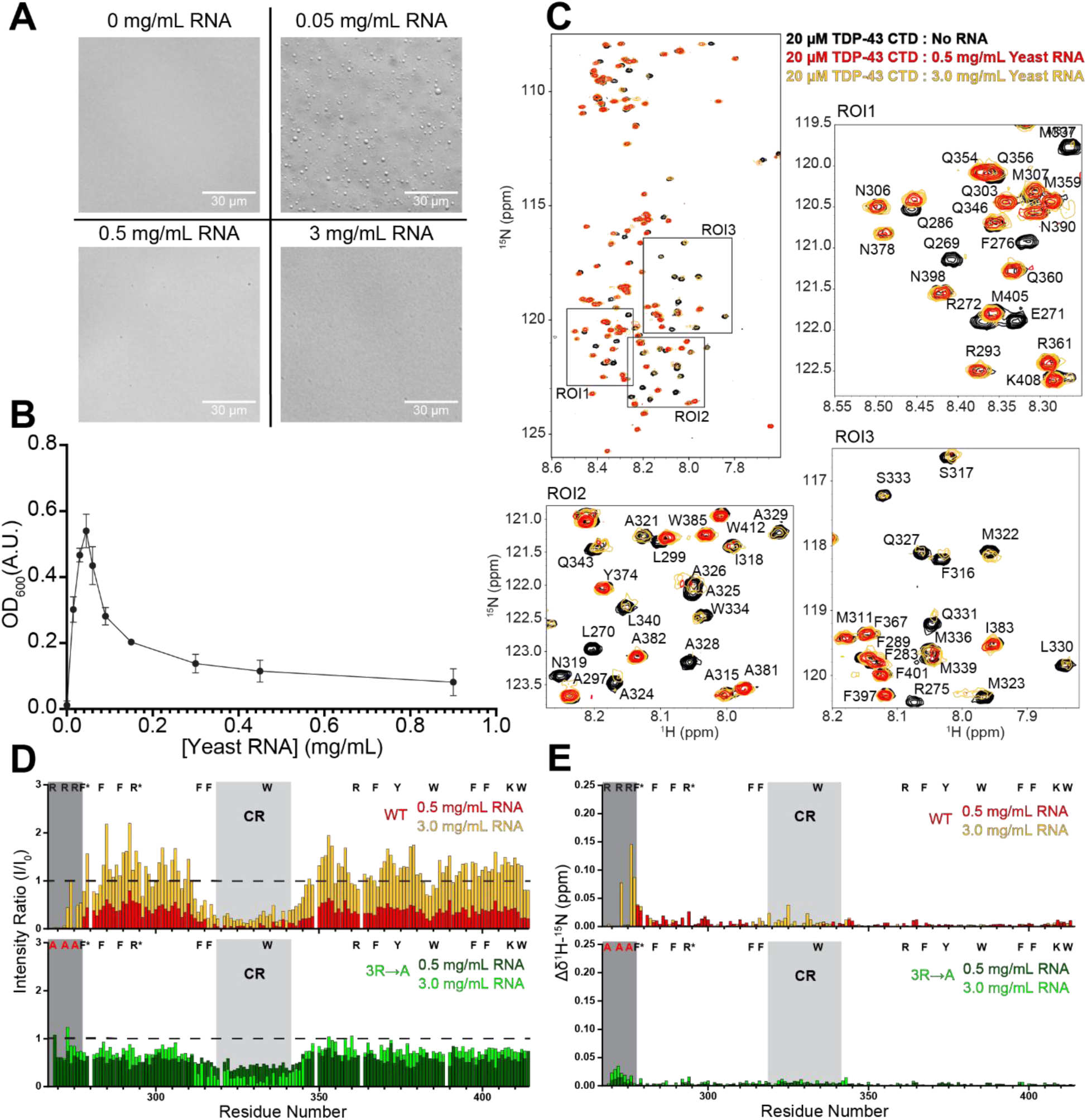
TDP-43 CTD Binds RNA Primarily Through a Positively Charged N-terminal Binding Region: **(A)** DIC micrographs of 20 μM WT TDP-43 CTD supplemented with 0.05 mg/mL, 0.5 mg/mL, and 3 mg/mL of Torula Yeast RNA (20 mM MES pH 6.1, 160 mM Urea) displaying reduction in protein droplets at higher RNA concentrations. **(B)** Turbidity assay measuring phase separation as 600 nm optical density of 20 μM WT TDP-43 CTD as a function of Yeast RNA concentration (20 mM MES pH 6.1, 160 mM Urea) demonstrating phase separation first peaks then decreases with increasing RNA concentration. **(C)** ^1^H-^15^N HSQC overlay of 20 μM WT TDP43-CTD (20 mM MES pH 6.1) with enlarged regions of interest (ROIs) in the absence (black) and presence of 0.5 mg/mL Torula Yeast RNA (red) and 3.0 mg/ mL Torula Yeast RNA (gold) displaying the loss of peak intensity at the N-terminus and conserved helical region due to RNA binding. **(D)** Normalized NMR peak intensity ratios comparing the change in intensity of 20 μM WT TDP-43 CTD and TDP-43 CTD 3RtoA titrated with 0.5 mg/mL Torula Yeast RNA and 3.0 mg/mL Torula Yeast RNA demonstrating that removal of N-terminal arginines disrupt RNA interactions. Relative positions of positively charged or aromatic residues are displayed above with RGG/FGG positions starred. The dark grey region corresponds to the N-terminal binding region lost upon addition of RNA while the light grey region displays the evolutionarily conserved region **(E)** Combined chemical shift perturbation plots of 20 μM WT TDP-43 CTD and TDP-43 CTD 3R→A titrated with 0.5 mg/mL Torula Yeast RNA and 3.0 mg/mL Torula Yeast RNA demonstrating significantly weakened CSPs after removal of arginines.

This observation allowed us to use NMR spectroscopy to investigate TDP-43 CTD interactions with RNA by acquiring spectra of the protein both before the addition of RNA and after addition of large amounts of RNA within the monophasic re-entrant condition. NMR fingerprint spectra (^1^H-^15^N heteronuclear single quantum coherence, HSQC) of TDP-43 CTD with 0.5 mg/mL and 3 mg/mL yeast RNA displayed the return of NMR signals (**Fig. 1C**) completely lost at lower concentrations of RNA (0.05 mg/mL). Consistent with turbidity measurements showing that complete re-entrance is not observed at 0.5 mg/mL RNA, NMR signal intensities for the re-entrant protein increase at 3.0 mg/mL compared to 0.5 mg/mL. Interestingly, when normalized to the intensity of the spectrum without RNA, many residues displayed a normalized intensity ratio greater than 1.0 for the 3.0 mg/mL titration point. In other words, many resonances show higher resonance intensity when (presumably) bound to excess RNA in the reentrant phase than in the absence of RNA (**Fig. 1D**). This result is atypical for NMR titrations where binding usually decreases signal intensity due to a combination of slowed reorientational motions or bound/unbound exchange that can both enhance NMR signal relaxation and attenuate resonances (89). Our data suggest that the binding of RNA can decrease conformational exchange and possibly other forms of exchange (**Fig. S1**), and that TDP-43 CTD variants with significantly decreased or abolished affinity for RNA (see below) do not display the same increase in intensity observed for the WT **(Fig 1D, Fig S2**).

Although addition of high (3.0 mg/mL) RNA results in many resonances either increasing in intensity or remaining similar to the condition without RNA, patches of other residues remained substantially decreased in intensity. These resonances lie in two distinct regions – the N-terminal region of the CTD and a highly evolutionarily conserved region (CR) in the center of the CTD previously characterized to form a α-helical structure critical for CTD multimerization (38–40,43,46). At the highest RNA concentrations, some attenuated N-terminal residue signals returned and displayed the greatest chemical shift perturbations (CSPs) (**Fig. 1E**). The observations that the resonances arising from the N-terminal region have large attenuation and CSPs are together strong indications that the N-terminal region acts as a direct RNA binding region. Given that this N-terminal region contained three (R268, R272, and R275) of the six positively charged residues present within TDP-43 CTD, we hypothesized that this region binds to RNA via ionic interactions with the negatively charged phosphate backbone. To confirm this, we created a variant with the three arginine residues mutated to alanine (R268A/R272A/R275A) which we refer to as TDP-43 CTD 3R→A. Unlike the WT, this variant does not strongly phase separate in the presence of RNA (**Fig. S2A**) and does not show attenuation of resonances for N-terminal residues (**Fig. S2B-D**), although we did observe small CSPs only near the N-terminus, which remains as a positively charged site due to the N-terminal amine (**Fig. 1E**). Curiously, at 3 mg/mL yeast RNA the CR of TDP-43 CTD still displays a reduction in resonance intensities, which we examine extensively (see below). These results demonstrate that while some interactions do still occur in the absence of the three N-terminal arginine residues, loss of these residues significantly disrupts CTD-RNA interactions, suggesting this N-terminal region serves as the primary but not only RNA binding site within CTD.

To further probe the residue specificity of the binding primary interaction, we created additional variants with the N-terminal arginines mutated to lysine (TDP-43 CTD 3R→K) and all arginine residues and the one lysine residue mutated to alanine (TDP-43 CTD allR/K→A) (**Fig. S2**). We found that our 3R→K variant behaved similarly to the WT suggesting that positive charge is required to facilitate the RNA interaction, but it is not arginine specific. Similarly, a variant that changes the two other TDP-43 CTD arginines and single lysine to alanine (TDP-43 CTD 2R/K→A) has a much smaller effect on phase separation than the N-terminal 3R→A variant, further suggesting the N-terminal arginines play a primary role in RNA interaction. The allR/K→A variant displayed no phase separation in the presence of RNA (**Fig. S2A**) and appeared not to interact with RNA at all based on near identical NMR spectra in the absence and presence of 0.5 mg/mL yeast RNA (**Fig. S2B-D**). Given the importance of ionic interaction for RNA binding, we also investigated how buffers of different ionic strengths by addition of either sodium chloride (NaCl) or a sodium phosphate buffer. Re-entrant behavior still occurred but required more RNA in these buffers with increased ionic strength, presumably due to screening of electrostatic interactions in solution (**Fig. S3A-B**). Importantly, NMR spectra using TDP-43 CTD A326P, a variant that discourages helix-mediated assembly resulting in superior NMR spectra in the sodium phosphate buffer condition, revealed that the primary features we observed above are present and hence independent of ionic strength – the N-terminal region still displays decreased intensity and strong CSPs upon addition of high amounts of RNA, which still induces peak intensity increases for some residues outside the CR (**Fig. S3C-F**).

### AAMD Simulations Reveal RNA Interactions Change within the Condensed Phase

To gain further molecular insights into CTD-RNA interactions and RNA-induced CTD conformational changes, we performed multi-microsecond all-atom molecular dynamics (AAMD) simulations of TDP-43 CTD in the presence of AUG12 RNA, a 12-nucleotide, unstructured sequence. AUG12 has high affinity to TDP-43 RRM domains and is representative of physiological TDP-43 targets (35,90,91) making it a suitable model system. We placed 10 RNA molecules around a single TDP-43 CTD chain and simulated the system for 12 µs (3 replicates x 4 µs each) to capture spontaneous binding events. Consistent with our hypothesis based on ^15^N-H chemical shifts, the N-terminal region engaged RNA primarily through arginine residues (R268/272/275), with additional contributions from neighboring polar (Q269) and aromatic (F276) residues as shown by replica-averaged contact probabilities (**Figure 2A**). Further, salt-bridge analysis between arginine nitrogen atoms and RNA backbone phosphate oxygens revealed persistent ionic interactions, with mean contact occupancies of 60% for R268, 45% for R272, and 66% for R275 across trajectories (**Figure 2B**). Notably, RNA interactions were much lower in the hydrophobic CR (**Figure 2A**), which maintained its α-helical structure (**Figure S4A**). In addition to the N-terminal interactions, we observed notable contacts in the C-terminal flanking region involving R361, F367, and W385/W412. We further probed the effect of RNA binding on the TDP-43 CTD conformational ensemble. We observed that RNA binding reduced intramolecular contacts between N- and C-terminal regions (**Figure S4B**), producing a slightly more expanded ensemble compared to the RNA-free state (**Figure S4C-E**).

**Figure 2:**
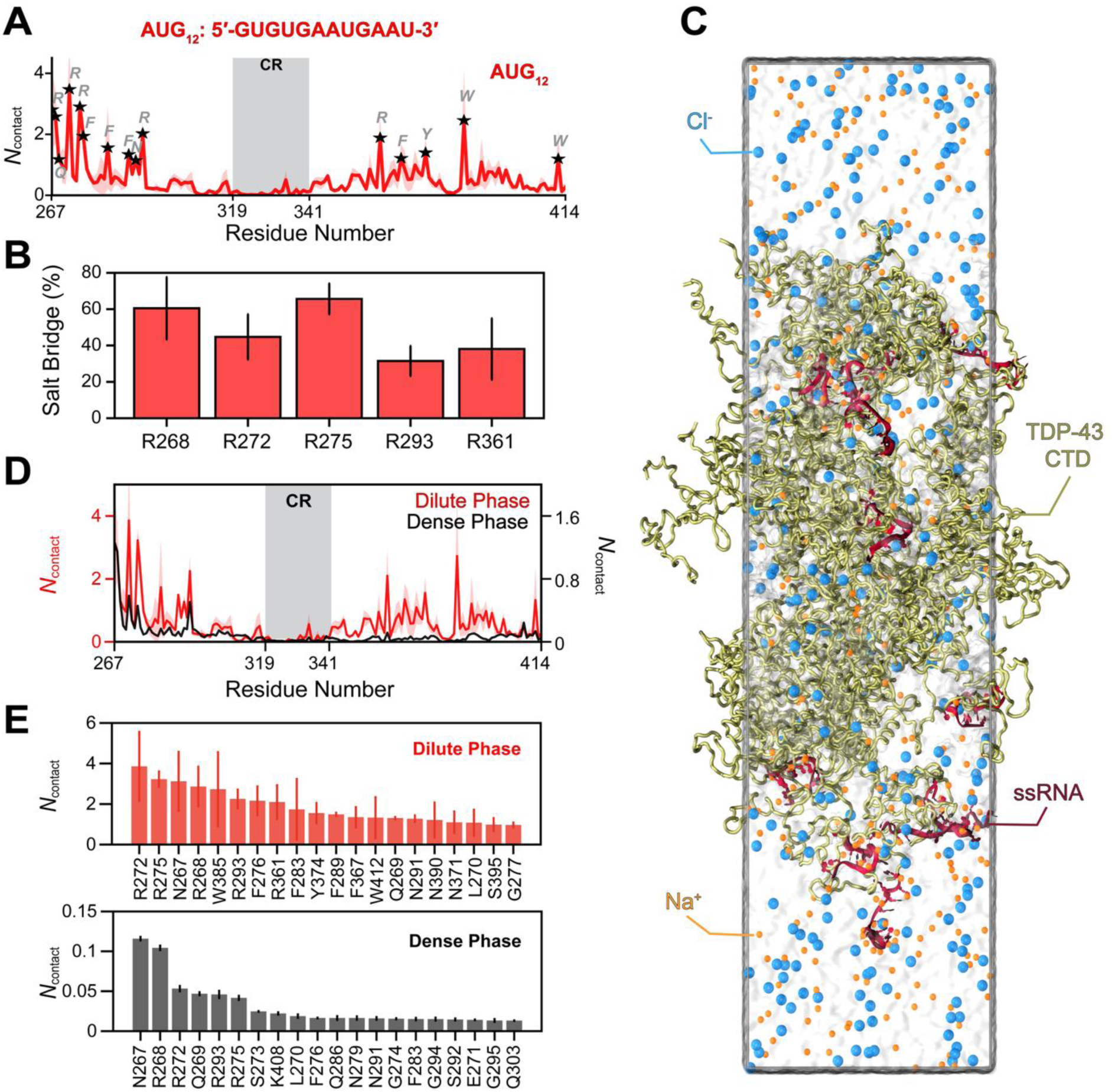
AAMD Simulations Reveal Differences in CTD-RNA Interactions in the Dense and Dilute Phase: **(A)** Protein-RNA contact profile along the CTD sequence. Contacts per protein residue summed across all 12 RNA nucleotide positions for AUG12 (5′-GUGUGAAUGAAU-3′). Contacts were calculated by counting heavy atom pairs within 4.5 Å cutoff distance for each trajectory frame, averaged across 10 RNA molecules and all frames. Data represent the mean with shaded regions indicating standard error of mean (SEM) from three independent simulations. Residues showing higher contact propensity are labeled with stars. **(B)** Percentage of salt bridge formation between arginine sidechains (R268: 60.50%, R272: 44.72%, R275: 65.62%, R293: 31.49%, and R361: 38.06%) and RNA phosphate groups. Salt bridges were identified in a binary manner across three independent simulations: if any sidechain nitrogen (NE, NH1, NH2) and any phosphate oxygen (O1P, O2P) were within 4 Å, a salt bridge was considered formed for that frame. Data show mean percentages ± standard error of the mean. **(C)** Representative snapshot from all-atom MD simulation (1 µs) containing 40 CTD chains and 11 RNA chains. **(D)** Comparison of protein-RNA contact profiles across the CTD sequence in dilute (red) and dense (black) phases displaying contacts per residue summed across all 12 nucleotide positions of AUG12 (5′-GUGUGAAUGAAU-3′). Contacts were defined as heavy atom pairs within 4.5 Å. For the dilute phase, data were averaged across 10 RNA molecules and all frames. For the dense phase, data were averaged across all CTD-RNA pairs and normalized by chain numbers and frame number. Shaded regions indicate standard error of the mean (SEM) from three independent simulations (dilute) or three trajectory blocks (dense). **(E)** Top 20 residues contributing most to RNA interactions and their corresponding in the dense (top) and dilute (bottom) phases.

Furthermore, to characterize the CTD-RNA interactions in the condensed phase, we performed phase-coexistence all-atom simulations of TDP-43 condensates (40 chains) based on a slab geometry with RNA (11 RNA molecules; **Figure 2C**) and without RNA. Notably, RNA interactions with the N-terminal arginine cluster in the dense phase were similar to those observed in the dilute phase (**Figure 2D-E).** In contrast, aromatic residues (F, Y, W) in the C-terminal region of CTD formed minimal contacts with RNA, suggesting that these interactions involving aromatic residues among CTD molecules outcompete RNA binding at these sites within the condensate. Notable differences in protein-RNA interactions between dilute and dense phases was also recently observed for FUS disordered domains in the presence of RNA (92). These findings imply that competing inter-protein interactions in dense phase can effectively modulate the protein-RNA interaction landscape. These observations are in striking contrast to homotypic prion-like condensates wherein the residue-level interactions formed in the dilute (i.e., intramolecular) and dense phases tend to be highly correlated (40,93).

Analysis of secondary structure further showed that CR helicity remained intact in the presence of RNA (**Figure S5A)**, and CR-CR inter-chain interactions were comparable in both condensates (**Figure S5B)**. Together, our simulations of the dilute and dense phases along with NMR experiments, support a model in which the arginines in CTD’s N-terminal segment serve as the primary interaction hub for RNA via electrostatic interactions with the phosphodiesters in the RNA backbone, while CR helicity and CR-mediated inter-chain interactions remain largely unaffected.

### RNA Induces TDP-43 CTD Multimer Formation by Acting as a Molecular Scaffold

Previous NMR studies of the CR of TDP-43 CTD demonstrate that residues 321-330 form a short α-helical structure which extends further to residues 331-343 following self-interactions that promote the formation of helical assemblies and TDP-43 CTD multimers (**Fig. 3A**) (38–42,46). With the exception of the allR/K→A variant, titration of CTD constructs with yeast RNA consistently resulted in reduction in signal of CR residues. However, the CR does not have any charged residues, making it unlikely to mediate direct contacts with RNA. In support of this argument, our AAMD simulations also showed no significant contacts between the CR and RNA in both dilute and condensed phases. Furthermore, CR helicity and CR-CR intermolecular interactions were preserved in the dense phase, regardless of the presence of RNA. Given the CR’s ability to form helical assemblies and these results, we hypothesized that the loss of NMR signals may be due to RNA-induced multimerization of the CR of several copies of TDP-43 CTD bound to the same long RNA, rather than direct interactions between the CR and RNA (**Fig. 3A**).

**Figure 3:**
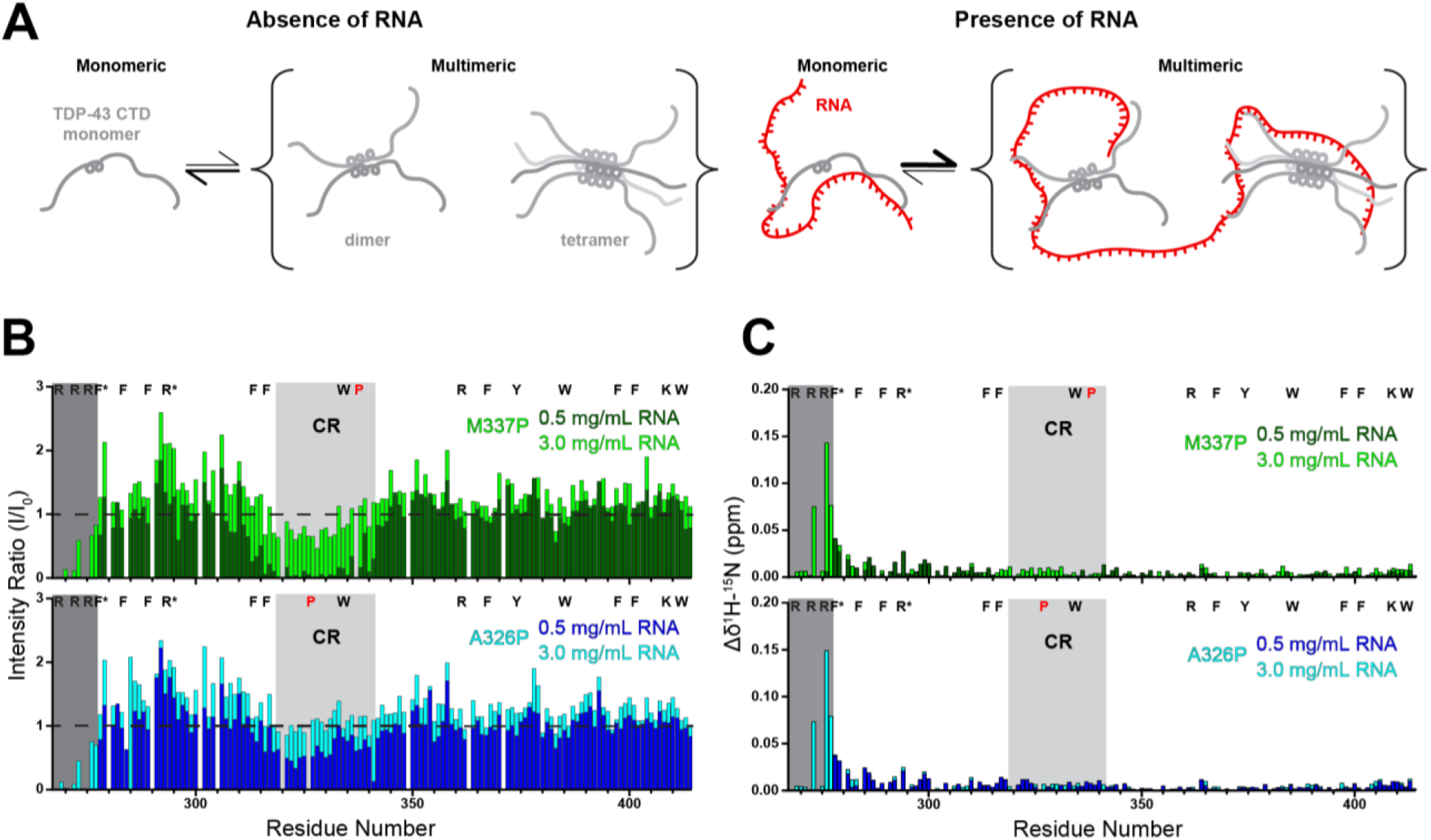
RNA Promotes TDP-43 CTD Protein-Protein Interactions and LLPS by Serving as a Molecular Scaffold: **(A)** Cartoon representation of proposed TDP-43 CTD multimerization via helical assemblies displaying our hypothesis that RNA promotes assembly. **(B)** Normalized NMR peak intensity ratios as function of residue position comparing the change in intensity of 20 μM TDP-43 CTD M337P titrated with 0.5 mg/mL Torula Yeast RNA (dark green) and 3.0 mg/mL Torula Yeast RNA (light green), and 20 μM TDP-43 CTD A326P titrated with 0.5 mg/mL Torula Yeast RNA (dark blue) and 3.0 mg/mL Torula Yeast RNA (light blue) demonstrating that disruption of helical structure prevents loss of conserved region residues especially at higher RNA concentrations. **(C)** Combined ^1^H -^15^N NMR CSPs of 20 μM TDP-43 CTD M337P titrated with 0.5 mg/mL Torula Yeast RNA (dark green) and 3.0 mg/mL Torula Yeast RNA (light green), and 20 μM TDP-43 CTD A326P titrated with 0.5 mg/mL Torula Yeast RNA (dark blue) and 3.0 mg/mL Torula Yeast RNA (light blue) demonstrating that N-terminal CSPs are preserved with mutation in conserved region.

To test this hypothesis, we tested how variants that decrease self-assembly of the CR affect the NMR resonances during RNA titrations. It is well established that TDP-43 CTD self-interactions depend on the helicity of the CR based on the behavior of variants that diminish or reenforce the helicity of the region (46). Two such helix-discouraging variants, TDP-43 CTD A326P and TDP-43 CTD M337P, diminish self-interactions by directly disrupting the formation of the α-helical structure or by impairing the extension and stabilization of the helix during the formation of cooperative helical multimers, respectively (38,46). NMR spectra of A326P and M337P titrated with 0.5 mg/mL and 3.0 mg/mL yeast RNA revealed loss of signal and/or dramatic shifting of N-terminal region resonances nearly identical to that of the WT, demonstrating that the N-terminal binding region is mostly unaffected by mutation of the CR (**Fig. 3B-C**). However, when titrated with RNA, the NMR resonances intensities from the conserved region entirely returned for A326P and partially returned for M337P (**Fig. 3B**), in stark contrast to the highly attenuated CR resonances observed for the WT. Compared to the large CSPs for the N-terminal region, these CR resonances showed no apparent CSPs in the conserved region (**Fig. 3C**). These results show that the observed signal attenuation in the CR requires CR assembly, further suggesting that RNA induces cooperative helical multimerization. We note that while the evidence here suggests RNA does not interact directly with the hydrophobic, uncharged CR, we cannot rule out that the multimeric CR creates an interaction site for RNA, though with no positively charged residues in the vicinity we do not think this is likely.

### Homopolymers Reveal Base Specific Differences in TDP-43 CTD RNA Interactions

Up until this point, we had limited our studies to using yeast RNA extract, as its sequence diversity allowed us to create a general model of TDP-43 CTD RNA interactions. However, it prevented us from probing the RNA sequence or structural specificity of the interactions. To investigate nucleotide base-specific contributions to CTD-RNA interactions we repeated our experiments with commercially available, synthetic, long (with an average size of approximately 800 kDa or ∼3000 nucleotides in length) RNA homopolymers (polyU, polyC, polyA, and polyG). Although these RNAs are much larger in size than the protein (about 50 times the mass), this is similar to the size of RNA transcripts and so provides a useful comparison where multiple proteins can simultaneously bind a single RNA. Measuring the turbidity of the CTD with each homopolymer revealed that while the RNA concentration at which the maximal turbidity was observed remained similar, there were base-specific differences in the reentrant behavior (**Fig. S6A**). We observed re-entrant behavior (loss of turbidity) with lower concentrations of polyG than the other homopolymers. Re-entrance with polyA was most distinct from all homopolymers, plateauing at an intermediate turbidity value and never fully reentering even at high RNA concentrations. Collecting NMR spectra of the CTD in the presence of extremely high concentrations (4.8 mg/mL) of each homopolymer also revealed profound differences in the overall signal intensities. Interestingly, the NMR spectrum of the CTD in the presence of 4.8 mg/mL polyG RNA resulted in almost no signals despite our turbidity measurements displaying no phase separation (**Fig. S6B-C**), suggesting that the CTD is strongly bound to RNA. Conversely, the CTD in the presence of 4.8 mg/mL polyU RNA had significantly higher overall peak intensity (except for the CR region) even when compared to the spectrum without RNA. The spectra collected in the presence of 4.8 mg/mL polyA and polyC had more comparable, weak resonance intensities overall, although the CR was far more attenuated with polyC than polyA. We also observed differences in the behavior of the N-terminal binding region as N-terminal residues decreased in peak intensity in the presence polyC and polyA RNA but underwent dramatic CSPs in the presence of polyU RNA (**Fig. S4D**). These experiments reveal base-specific differences in the binding modes of CTD to homopolymer RNAs. However, the much larger size of the RNA compared to TDP-43 CTD and the variable distribution of sizes of the RNAs preclude elucidation of some of the specific interaction details, which we address below.

While our initial NMR experiments demonstrate differences in CTD RNA interactions depending on the nucleotide base, observing changes to the amide backbone environment alone did not provide a clear molecular basis for why these differences occur. To dissect the molecular determinants of base-specific interactions, we performed AAMD simulations of TDP-43 CTD monomer (dilute phase) in the presence of four distinct RNA homo-oligonucleotides (12mers: rG_12,_ rU_12,_ rA_12_, and rC_12_), accumulating 48 µs of simulation time (4 sequences x 3 replicates x 4 µs each) (**Fig. S7**). To quantify the molecular recognition patterns between CTD and RNA, we calculated the averaged contact frequencies between CTD sequence and each RNA homo-oligonucleotide (**Fig. 4A)**. All homo-oligonucleotides displayed a clear preference for arginine residues within the N-terminal flanking region, consistent with our previous NMR and MD observations, confirming this region serves as the primary interaction hub regardless of RNA sequence. Notably, rG_12_ and rU_12_ formed significantly more contacts across the CTD sequence, particularly with the C-terminus of CTD, compared to rA_12_ and rC_12_ (**Fig. S8**). Furthermore, CR region showed minimal interactions across all sequences, except rG_12_, which showed enhanced contacts with W334.

**Figure 4:**
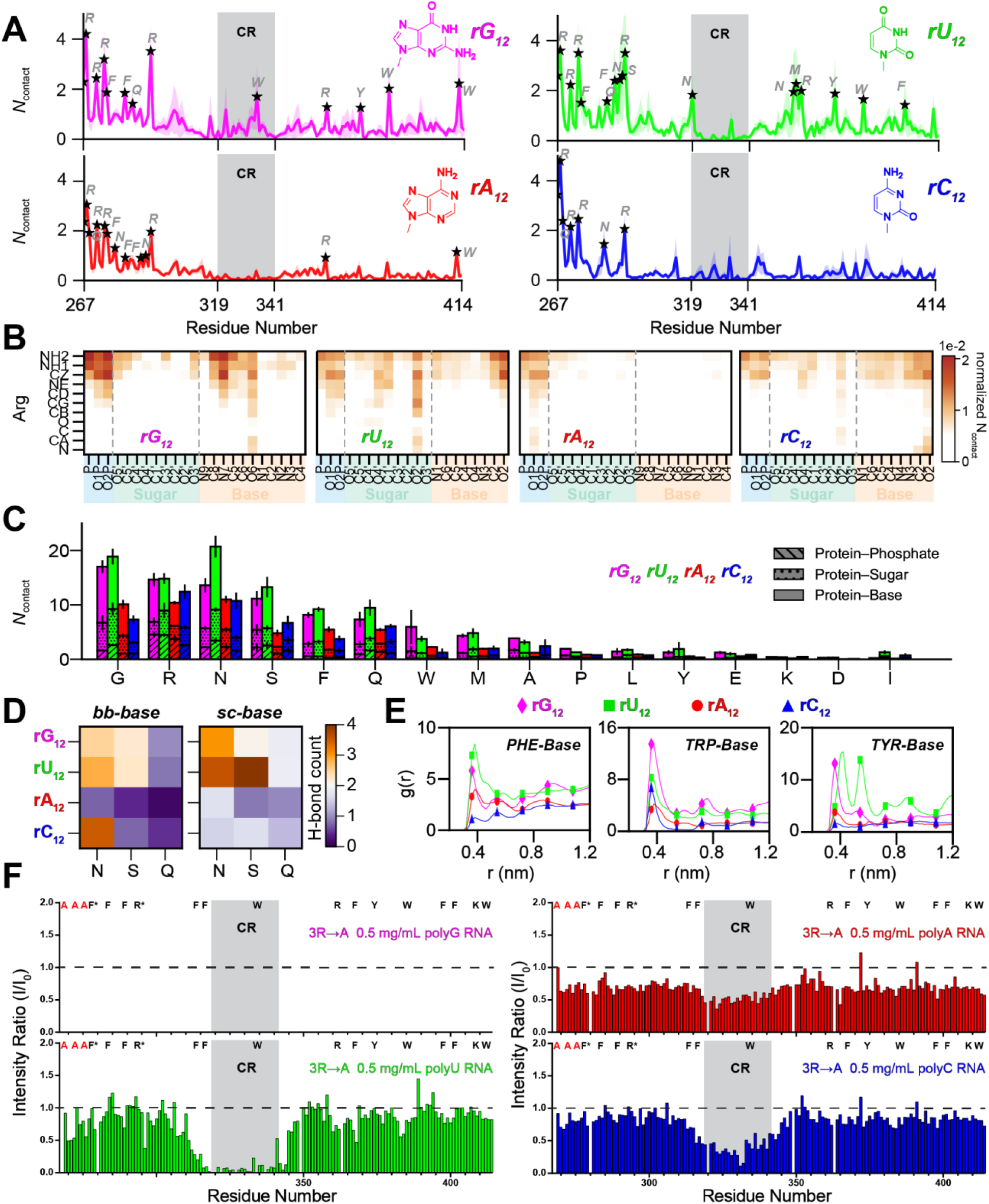
MD Simulations and Alanine Screens Demonstrate Greater Interaction Between CTD With U and G-Rich RNA Sequences: **(A)** Protein-RNA contact profile along the CTD sequence. Contacts per protein residue summed across all 12 RNA nucleotide positions for homo-oligonucleotide RNA sequences (rG_12_, magenta; rU_12_, green; rA_12_, red; rC_12_, blue). Contacts were calculated by counting heavy atom pairs within 4.5 Å cutoff distance for each trajectory frame, averaged across 10 RNA molecules and all frames.

Data represent the mean with shaded regions indicating standard error of mean (SEM) from three independent simulations. Residues showing higher contact propensity are labeled with stars. **(B)** Atomic contact map between heavy atoms of IDR1 Arginine residues (R268, R272, R275, R293) and heavy atoms of RNA nucleotides. Contact pairs were identified using a 4.5 Å cutoff and are normalized by 120 total nucleotides (10 RNA molecules × 12 nucleotides each), representing contacts per nucleotide per frame. Data show mean from three independent simulations. **(C)** Protein-RNA contact decomposition by amino acid type. Contacts summed across all residues of each amino acid type for homo-oligonucleotide RNA sequences (rG_12_, magenta; rU_12_, green; rA_12_, red; rC_12_, blue). Stacked bars show contributions from protein contacts with RNA phosphate, sugar, and base groups. Error bars indicate SEM from three replicate simulations. U- and G- bases have preference for polar (N, S, Q) and aromatic (F, W, Y) residues while all bases have enhanced interactions with arginine side chain potentially through electrostatic interactions (cation-π and/or hydrogen bonding). **(D)** Hydrogen bond analysis of polar residues with RNA bases. Hydrogen bond counts for asparagine (N), serine (S), and glutamine (Q) residues with homo-oligonucleotide RNA sequences (rG_12_, magenta; rU_12_, green; rA_12_, red; rC_12_, blue) are counted and shown as heatmap. Hydrogen bonds were identified by donor-acceptor distances ≤3.5 Å and decomposed into backbone-base (bb-base) and sidechain-base (sc-base) interactions. Values represent the sum of H-bonds between all residues of each type and any nucleotide (10 RNA molecules with 12 nucleotides each), averaged over trajectory frames. Data show mean from three independent simulations. **(E)** Radial distribution functions (RDF) between aromatic residues and RNA bases. g(r) plots showing the spatial distribution of RNA bases (10 RNA molecules with 12 nucleotides each) around aromatic residues PHE, TRP, and TYR for homo-oligonucleotide RNAs (rG_12_, magenta; rU_12_, green; rA_12_, red; rC_12_, blue). RDFs were calculated using the center of mass of aromatic ring atoms (PHE: CG, CD1, CD2, CE1, CE2, CZ; TRP: CD2, CE2, CZ2, CH2, CZ3, CE3; TYR: CG, CD1, CD2, CE1, CE2, CZ) and nucleotide base atoms (N1, C2, N3, C4, C5, C6). Peaks indicate preferential distances for π-stacking and hydrophobic interactions. Data averaged from three replicate simulations. **(F)** Normalized peak intensity ratio plots of 20 μM TDP-43 CTD 3RtoA in the presence of 0.5 mg/mL polyG RNA (magenta), polyU (green), polyA (red), and polyC RNA (blue) displaying conserved region peak intensity decreased the most with polyU RNA compared to polyC and polyA RNA and complete loss of polyG signal.

To explore how base identity affects interactions of the four N-terminal arginine residues (R268, R272, R275, R293), we constructed atomic-level contact maps to identify pairwise interactions between atom types of the arginine sidechain and RNA (**Figure. 4B)**. While ionic interactions with the phosphate backbone remained dominant across all sequences explaining this region’s charge-based affinity for RNA, rG_12_, rU_12_, and to a lesser extent rC_12_ formed additional base-specific contacts with arginine side chains. In addition to N-terminal arginines in the CTD, we further investigated whether other residue types could contribute to base-specific recognition. Our analysis indicated that glycine, polar (N, S, Q) and aromatic residues (F, W, Y) interacted more favorably with rG_12_ and rU_12_ bases compared to rA_12_ and rC_12_ bases (**Fig. 4C**). Glycine residues are abundant in CTD and proximal to arginine (R) and aromatic (F, W, Y) residues, explaining the enhanced contacts with RNA. Hydrogen bond analysis (donor-acceptor distance ≤ 0.35 nm) confirmed that G and U bases form more H-bonds with polar residues (N, S, Q) than A and C bases, predominantly between the sidechain and base moieties (**Fig. 4D**). To further characterize the aromatic interactions, we calculated radial distribution functions (RDFs) between aromatic ring centers (F, W, Y) and nucleotide bases. RDFs revealed sharp peaks at around 0.4 nm for rG_12_ and rU_12_, characteristic of π-stacking interactions (**Fig. 4E**). While G and U bases showed enhanced stacking with all three aromatic residues compared to other bases, G bases notably exhibited higher stacking propensity with Trp residues compared to U bases. Taken together, our results suggest that preferential recognition of G and U bases is facilitated through a network of electrostatic (ionic interactions and hydrogen bonding) and hydrophobic (aromatic stacking) interactions, enabling both the N-terminal and C-terminal flanking regions to selectively engage with G and U bases.

Building upon these findings and based on our model that CTD interactions with polyA and polyC are more dependent on the N-terminal binding region, we hypothesized that the removal of the N-terminal arginine residues should have a much greater impact on interactions with polyA and polyC compared to polyG and polyU. To test this hypothesis experimentally, we collected NMR spectra of TDP-43 3R→A with 0.5 mg/mL of each long-chain homopolymer. For polyU, polyA, and polyC we observed no loss of signal at the N-terminal binding site, as expected due to removal of the arginine residues. However, we did observe that resonances within the CR were completely attenuated in the presence of polyU RNA, partially attenuated in the presence of polyC, and only slightly attenuated in the presence of polyA (**Fig. 4F, Fig. S9A**). We also observed the magnitude of CSPs followed the same trend with polyU inducing the largest shifts and polyA the smallest (**Fig S9B)**. In the presence of polyG, similar to the WT TDP-43 CTD, all resonances in the 3R→A spectrum were attenuated below the noise, demonstrating not only the presence of interactions outside the N-terminal binding region but also that these interactions are much stronger with polyG than with the other homopolymers. These results demonstrate that polyU still strongly drives multimerization of the protein and retains a stronger interaction than polyA and polyC in the absence of the N-terminal arginines, lending support to our interaction model.

To determine if we could completely abolish RNA interactions with the homopolymers as we did with yeast RNA, we also collected NMR spectra of allR/K→A titrated with 0.5 mg/mL of each long homopolymer (**Fig. S9C-D**). While we observed nearly no change to the NMR spectra in the presence of 0.5 mg/mL of polyU, polyA, and polyC RNA, we observed decreases in peak intensity for the spectrum in the presence of 0.5 mg/mL polyG RNA. The residues undergoing the greatest change in intensity correspond to those in the N-terminal disordered region (IDR1) and the CR while signals of C-terminal disordered region (IDR2) residues were consistently higher. Therefore, interactions with polyU, polyC, and polyA are heavily dependent on charged residues, while polyG is uniquely able to interact with TDP-43 CTD via other modes. We hypothesize that these differences may arise due to the ability of polyG to form higher order structures that we explore below.

### TDP-43 CTD Binds Strongly to Quadruplexes Structures Via Aromatic Interactions

The homopolymer RNAs are all much larger than TDP-43 CTD, may not have precisely the same size distribution, and are large enough to serve as scaffolds for TDP-43 CR assembly, binding multiple copies of TDP-43 CTD. In order to better compare the base-specific details of the interactions with TDP-43 CTD, we decided to test TDP-43 CTD binding to short ssDNA 12mer homo-oligonucleotides. We created 12mers for each canonical base (dT_12_, dA_12_, dG_12_, dC_12_,) in addition to a noncanonical polyU ssDNA 12mer (dU_12_) to compare to the biological RNA base. NMR spectra of TDP-43 CTD in the presence of 180 μM homo-oligonucleotides (a 9-fold molar excess equivalent to ∼0.7 mg/mL) displayed two different interaction profiles, with dT_12_, dA_12_, and dU_12_ displaying large CSPs and relatively minor signal attenuation, while dG_12_ and dC_12_ displayed no CSPs but significant peak attenuation across the sequence (**Fig. 5A-B, Fig. S10A-B**). The largest CSPs for dT_12_, dA_12_, and dU_12_ were localized to the N-terminal binding region (**Fig. 5B**), with dT_12_ and dU_12_ displaying near identical CSPs (in both magnitude and direction), suggesting that the single methyl group difference between U and T does not affect these interactions. CSPs and resonance intensity ratios for dA_12_ differed from dT_12_ and dU_12_ in both magnitude and direction (i.e. amount/direction of CSP in ^1^H and ^15^N) (**Fig. 5A-B**), demonstrating the base can play a role in determining the chemical environment of the binding region. Importantly, dT_12_, dA_12_, and dU_12_ did not display significant reduction in CR signal intensity relative to the disordered regions, unlike the long homopolymers (**Fig. S10B**), suggesting short 12mers do not induce multimerization at these conditions with molar excess of the oligonucleotides. Conversely, spectra in the presence of dG_12_ and dC_12_ displayed significant signal attenuation of residues across the entirety of IDR1 and the CR (**Fig. 5A**). We also observed localized attenuation in IDR2 located around aromatic residues within the sequence, particularly near W385, suggesting aromatic residues may play a larger role in interactions with dG_1_

**Figure 5:**
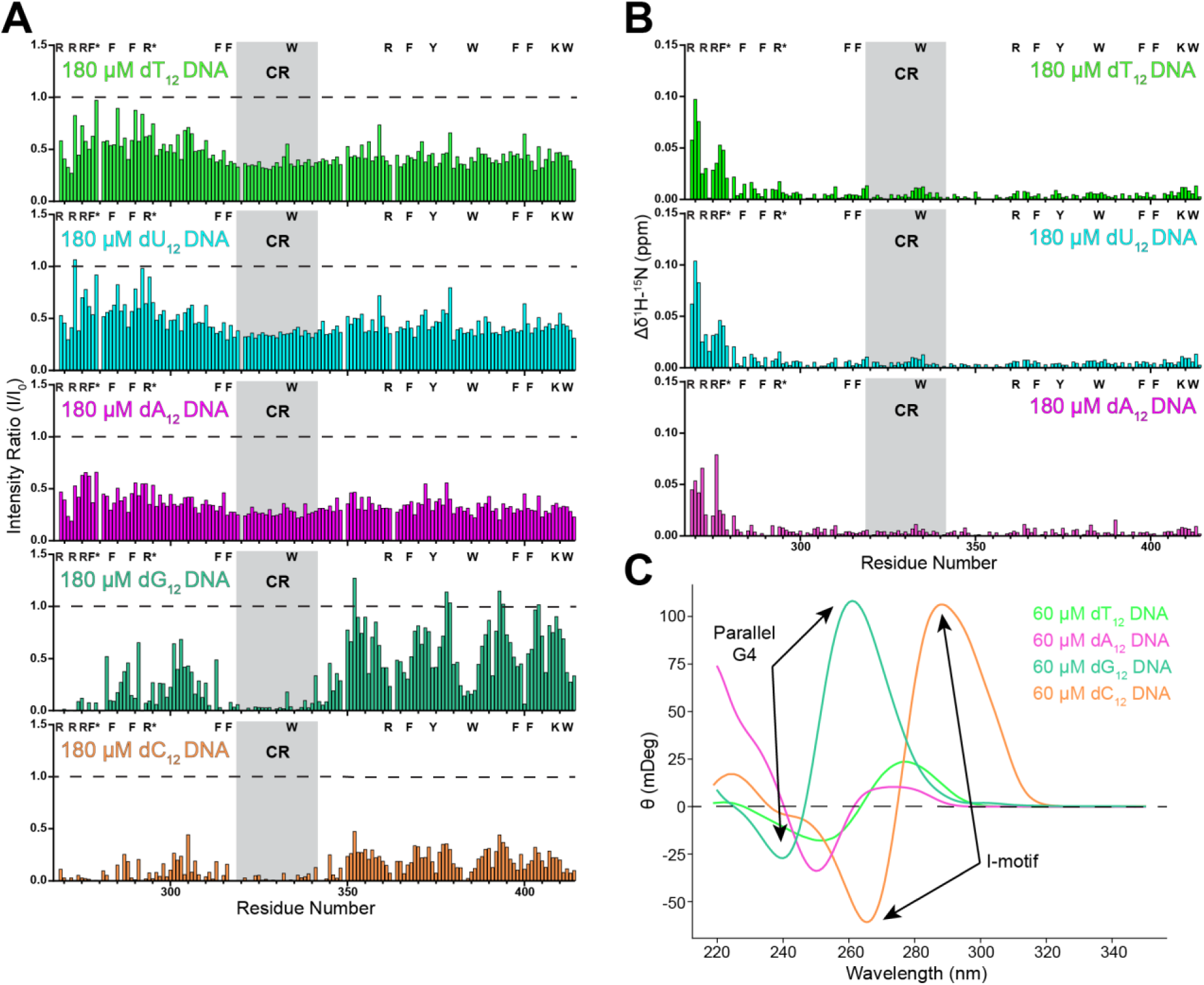
Small ssDNA Homo-oligonucleotides Reveal Structure Specific Differences in Binding Interaction: **(A)** Normalized NMR peak intensity ratios comparing the change in intensity of 20 μM WT TDP-43 CTD residues (20 mM MES, pH 6.1) in the presence of 180 μM dT_12_ (light green), 180 μM dU_12_ (light blue), 180 μM dA_12_ (magenta), 180 μM dG_12_ (ocean green) and 180 μM dC_12_ (orange) demonstrating the protein has two different interaction modes depending on homopolymer utilized. **(B)** Combined ^1^H -^15^N NMR chemical shift perturbations of 20 μM WT TDP-43 CTD residues (20 mM MES, pH 6.1) in the presence of 180 μM dT_12_ (light green), 180 μM dU_12_ (light blue), and 180 μM dA_12_ (magenta). dT_12_ and dU_12_ display comparable CSPs while dA_12_ displays difference in magnitude. **(C)** CD spectra overlay of 60 uM dT_12_ (light green), 60 uM dA_12_ (magenta), 60 uM dG_12_ (ocean green), and dC_12_ (orange) (20 mM MES, pH 6.1). dG_12_ displays a strong positive peak at 260 nm and a negative peak at 240 nm indicating the presence of parallel G4 quadruplex structure. dC_12_ displays a strong positive peak at 290 nm and a strong negative peak at 265 nm indicating the presence of an I-motif. dT_12_ and dA_12_ display less secondary structure based on smaller ellipticity values.

Given our observation that the interaction with dG_12_ and dC_12_ is distinctly different when compared to dT_12_, dA_12_, and dU_12_, this discrepancy could be due to the ability of G-rich RNA and DNA sequences to form G4 quadruplexes and C-rich DNA sequences (but not C-rich RNA sequences, due to interference by the 2’ OH in rC) to form I-motifs (94–96). Importantly, these higher order nucleic acid structures were not modeled in our simulations. Indeed, circular dichroism (CD) spectra of each canonical 12mer revealed that dG_12_ and dC_12_ displayed significantly higher CD absorption than dT_12_ and dA_12_ at the same concentration (**Fig. 5C**), demonstrating greater presence of secondary structure. Additionally, the CD spectrum of dG_12_ displayed the characteristic peaks of a parallel G4 quadruplex (97), while the CD spectrum of dC_12_ displayed the characteristic peaks of an I-motif (96). These data suggest that the profound difference in TDP-43 CTD interactions strongly depends on the presence of DNA secondary structure and supports previous reports of strong G4 quadruplex binding by the CTD (55–58). To provide additional evidence that this unique interaction mode is specific to the presence of structures rather just G-rich or C-rich sequences, we decided to characterize the interaction of TDP-43 CTD with a 34mer ssDNA oligo based on the c-Myc promoter sequence, previously demonstrated to form parallel G4-quadruplex structures and strongly bind to TDP-43 CTD (**Fig. 6A**) (57,94). We also characterized the interactions of the CTD with the reverse complement of this G4 sequence which had previously been demonstrated to be able to form an I-motif (**Fig. 6B**) (96). CD spectra collected of the ssDNA c-Myc sequences (**Fig. 6C**) were nearly identical to the spectra of the dG_12_ and dC_12_ homopolymers (**Fig. 5C**) confirming that despite possessing different sequences and lengths the c-Myc sequences produced similar higher-order structures to the homo-oligonucleotides (**Fig. 6C**).

**Figure 6:**
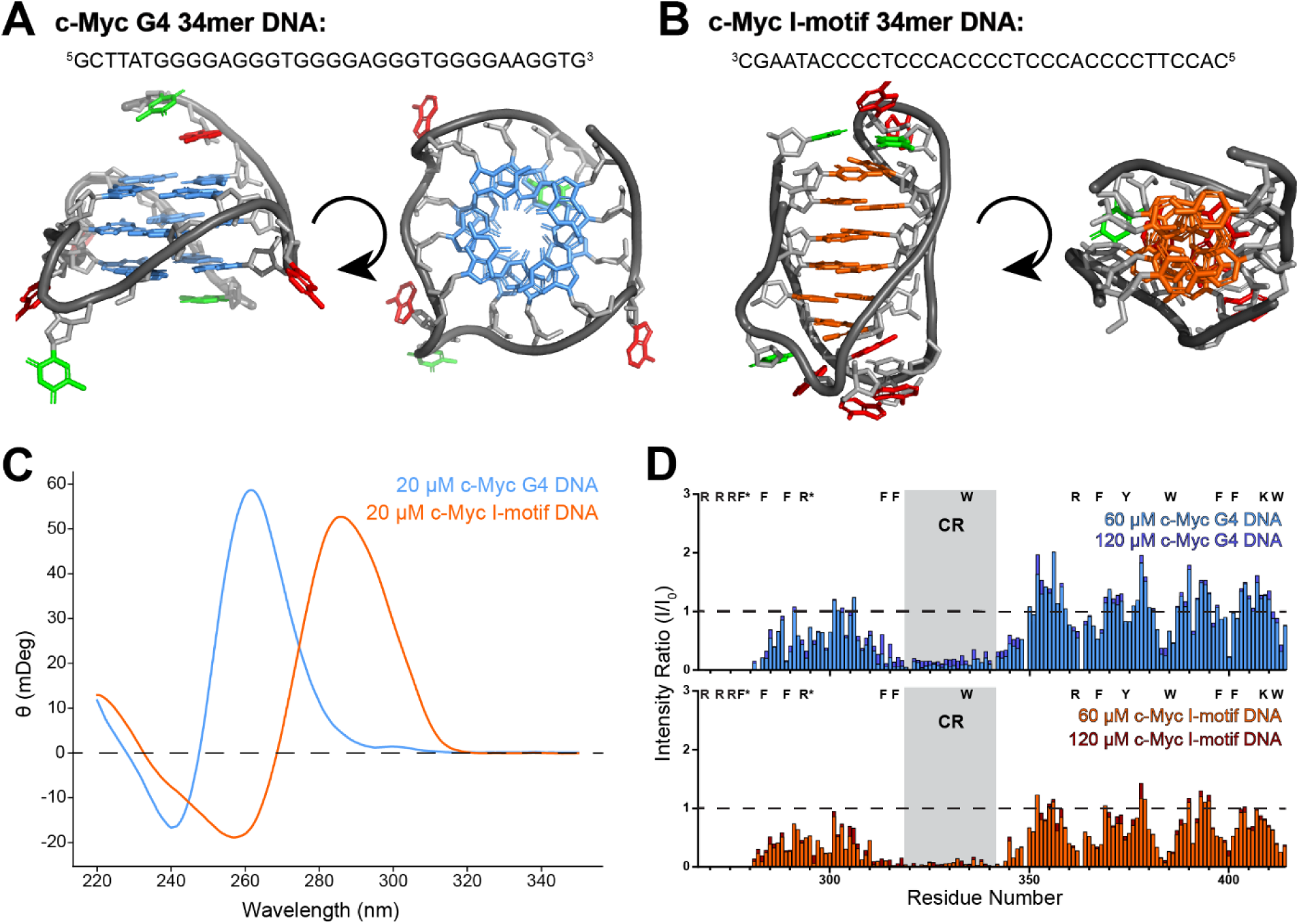
TDP-43 CTD Binds Strongly to Sequences Forming Parallel G4 Quadruplexes and I-motifs Via Aromatic Interactions: **(A)** Sequences of 34mer ssDNA sequences based on the c-Myc promotor known to form a G4 quadruplex and the published structure of the c-Myc promotor G4 quadruplex (PDB: 2LBY) (98). **(B)** Sequences of 34mer ssDNA sequences based on the c-Myc promotor known to form an intramolecular I-motif and the published structure of a structurally analogous telomere I-motif (PDB: 1EL2) (99). **(C)** CD spectra overlay of 20 uM c-Myc G4 quadruplex DNA (sky blue), and 20 uM c-Myc I-motif DNA (orange) (20 mM MES, pH 6.1). c-Myc DNA displays a strong positive peak at 260 nm and a negative peak at 240 nm indicating the presence of parallel G4 quadruplex structure while a strong positive peak at 290 nm and a strong negative peak at 265 nm indicating the presence of an I-motif. **(E)** Overlayed normalized NMR peak intensity ratios as a function of sequences position comparing the change in intensity of 20 μM WT TDP-43 CTD (20 mM MES, pH 6.1) titrated with 60 μM c-Myc G4 (sky blue) and 120 μM c-Myc G4 (purple) or 60 μM c-Myc I-motif (orange) and 120 μM c-Myc I-motif (dark red) displaying similar interaction to dC_12_ and dG_12_ homopolymers and little change in peak intensity with greater DNA concentration.

Probing the details of the interactions with these structured oligos, NMR spectra of TDP-43 CTD titrated with 60 μM and 120 μM concentrations of c-Myc G4 quadruplex or c-Myc I-motif revealed that the protein had a similar interaction with both constructs (**Fig. 6D**). Similar to dG_12_ and dC_12_, resonances in the presence of either quadruplex were significantly attenuated, with the most significant attenuation localized to the entire N-terminal binding region and the CR (**Fig. 6D, Fig. S10C**). We also once again observed dramatic attenuation of some IDR2 residues with both G4 quadruplexes and C-rich I-motifs that appeared to be localized around aromatic residues. Signals in the presence of the c-Myc I-motif were consistently lower compared to the c-Myc G4 quadruplex, suggesting the protein may bind the I-motif with greater affinity. Doubling the concentration of either sequence resulted in little change in resonance intensity for any residue, suggesting the interaction is saturated at the concentrations tested. Based on these results we concluded that the previously reported strong binding of TDP-43 CTD to G4 quadruplex structures is due to a greater contribution from aromatic residues to the binding interaction compared to unstructured RNA/DNA sequences. We also found that this interaction is not G4 quadruplex specific and that TDP-43 CTD binds strongly to I-motif structures by similar molecular mechanisms.

### TDP-43 CTD Displays Greater Complex Formation and Binding Affinity with TG-Rich Sequences Compared to AC-Rich Sequences

TDP-43 has been well documented to preferentially bind UG-rich RNA and TG-rich DNA sequences, however this strong binding affinity has been attributed mainly to its two folded RRMs, particularly RRM1 (34,35,91). Our NMR experiments and MD simulations reveal that the CTD seems to also possess a higher affinity for U and G-rich sequences over C and A-rich, although the high binding affinity for polyG sequences observed experimentally is affected by the presence of quadruplex structures that was not modeled in the simulations. Given that TG-rich and AC-rich DNA sequences are incapable of forming quadruplex structures due to their alternating base sequence, we tested if there were any detectable differences in CTD binding in the absence of RNA/DNA structure, and how this compared to the binding preference of the RRMs. To accomplish this, we created two new ssDNA 34mer oligonucleotides composed of 17 TG or AC repeats which we refer to as (TG)_17_ and (AC)_17_, respectively (**Fig. 7A**). NMR spectra of TDP-43 CTD with the 34mers revealed that both (TG)_17_ and (AC)_17_ induced strong CSPs in the N-terminal region, with the direction and magnitude of these chemical shifts being largely similar, consistent with the binding of these sites to RNAs regardless of the base identity, as we showed above (**Fig. 7B, Fig. S11A-B**). These N-terminal CSPs remained effectively the same at every concentration tested, demonstrating that the N-terminal binding region is saturated within the reentrant state at these conditions. However, at the two lower concentrations, CSPs within the CR were greater in the presence of (TG)_17_ than (AC)_17_, consistent with induction of more CR-CR contacts by (TG)_17._ At the highest concentrations where the oligos are in high excess, both (TG)_17_ and (AC)_17_ had significantly lower CSPs, most likely due to helix-helix interactions being diminished when each copy of TDP-43 binds its own oligo. Importantly, when we compare the average peak intensity of each structural region of the protein, we also observed differences between the two ssDNA sequences, (**Fig. 7C, Fig. S11C**), with the average intensity of the CR being consistently lower in the presence of (TG)_17_ when compared to the same concentration of (AC)_17_, suggesting greater interaction with (TG)_17_. At the two highest oligo concentrations, we also observed nearly no change in the average peak intensity of the IDRs with (TG)_17_ while the IDRs show higher intensities at higher concentrations of (AC)_17_, suggesting that saturating the interactions requires more (AC)_17_, again suggesting stronger interaction with (TG)_17_.

**Figure 7:**
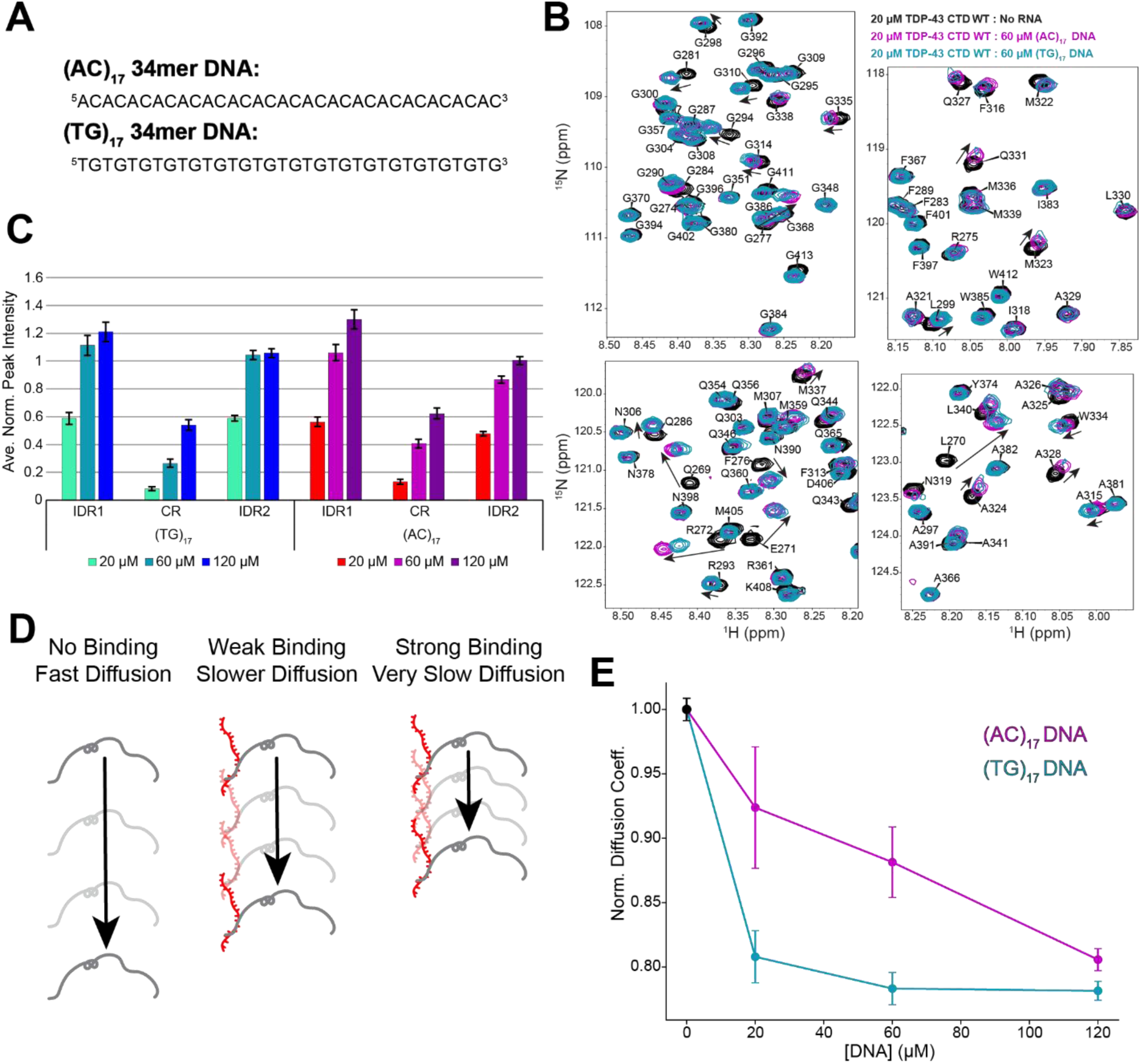
TDP-43 CTD RNA Interactions Achieve Saturation Faster with TG-Rich Sequences Compared to AC-Rich: **(A)** Sequences of (TG)_17_ and (AC)_17_ ssDNA 34mer constructs used to test sequence affinity. **(B)** ^1^H-^15^N HSQC ROIs of 20 μM WT TDP43-CTD (20 mM MES pH 6.1) in the absence (black) and presence of 60 uM (TG)_17_ DNA (light blue) and 60 uM (AC)_17_ DNA (magenta) displaying similar N-terminal CSPs for both constructs but differences in conserved region CSPs and intensities. **(C)** Average normalized peak intensity of each region of the CTD as function of (TG)_17_ or (AC)_17_ concentration displaying differences in the intensities of each region depending on ssDNA concentration and sequence. Error bars represent the standard error of the mean. **(D)** Cartoon demonstrating how binding interactions of varying strength will impact the average rate of diffusion of a protein and our hypothesis that stronger (TG)_17_ binding will result in a slower diffusion rate compared to (AC)_17_. **(E)** Plot of the apparent diffusion coefficient of TDP-43 CTD, normalized to the diffusion coefficient in the absence of DNA (black), as a function of increasing (AC)_17_ (magenta) and (TG)_17_ (light blue) concentration displaying the protein reaches a point of saturation faster with (TG)_17_ than (AC)_17_.

Because phase separation upon addition of substoichiometric amounts of nucleic acid oligos precludes direct measurement of binding affinity, to provide additional evidence of stronger CTD binding to (TG)_17_ compared to (AC)_17_, we measured the apparent diffusion coefficient of TDP-43 CTD when titrated with (TG)_17_ and (AC)_17_ using Diffusion-Order NMR Spectroscopy (DOSY-NMR). The diffusion coefficient of a molecule through solution is directly correlated to its hydrodynamic radius. As a result, binding events that result in the formations of a much larger complex, such as a protein bound to a DNA oligonucleotide, result in a detectable change in the protein’s diffusion coefficient. Stronger binding events in turn result in greater changes to the diffusion coefficient as the protein spends more time in the larger slower moving complex compared to a weaker binder (**Fig. 7D**). When we measured the diffusion coefficient of TDP-43 CTD titrated with either (TG)_17_ or (AC)_17_ then normalized to the diffusion coefficient in the absence of DNA (**Fig. 7E**), we observed that the apparent diffusion coefficient of the CTD decreased in the presence of both ssDNA sequences, but consistently displayed a lower diffusion coefficient in the presence of (TG)_17_ when compared to the same concentration of (AC)_17_ despite only a marginal (5.1% higher) difference in the mass of (TG)_17_ oligo. Additionally, we observed that while diffusion of the protein decreased consistently at every (AC)_17_ concentration tested, the diffusion coefficient appeared to plateau with addition of stoichiometric amount of (TG)_17_, providing additional evidence the binding interaction saturates faster with (TG)_17_ compared to (AC)_17_. Together these results suggest that TDP-43 CTD’s preference for U and G containing RNA sequences may cooperatively assist in the binding of UG-rich RNAs to the TDP-43 RRMs.

## Discussion

Here, we present our findings on the molecular-level interactions between the largely disordered CTD of TDP-43 and RNA to uncover how RNA sequence and structure can give rise to preferential interactions with a prion-like domain. We demonstrate that TDP-43 CTD contains a non-specific disordered RNA-binding region spanning residues 268-279, characterized by three positively charged arginine residues clustered within this region. At lower RNA concentrations, where the molar concentration of large RNAs is much lower than that of TDP-43 CTD, RNA appears to strongly induce the formation of helical assemblies between different CTD monomers. In this context, RNA serves as a molecular scaffold, creating a locally high protein concentration around individual RNAs promoting greater protein-protein interactions. Reducing the affinity of the protein for RNA by removing the positively charged residues from the N-terminal binding region reduces both the ability of RNA to promote phase separation and the formation of these multimers via protein-protein helical contacts, suggesting that the ability of RNA to promote TDP-43 phase separation is somewhat coupled to its scaffolding ability (100,101).

Despite lacking a classical RNA binding domain, the CTD exhibits some degree of base specificity, displaying a preference for binding U- and G-rich sequences over A- and C-rich sequences. The molecular basis for this base selectivity may lie in the chemical properties of the nucleobases. When comparing the ring structures, we noticed that U and G possess an additional ketone group when compared to their purine or pyrimidine counterpart, suggesting this substituent may play a role in defining the interaction. Indeed, our simulations suggest that the guanidino group of arginine appears to interact with the ketone groups of G, U, and C. Additionally, we noticed that U and G share a common structural feature in that they possess a protonated aromatic nitrogen directly next their ketone groups, which differs from C that contains a ketone but lacks a protonated nitrogen, and A which lacks both of these features. This difference could account for the stronger interactions between U and G with polar residues observed as this combination of a hydrogen-bond accepting ketone and hydrogen-bond donating protonated nitrogen could interact more favorably with the amide or hydroxyl group of polar residues. We also observed that the multiple FG(G) and WG motifs present within the CTD appeared to preferentially form hydrophobic stacking interactions with the U and G nucleotide rings compared to C and A, further driving base-specificity. Future studies using modified bases or synthetic base analogs (102) could be utilized to further probe how ring substituents and protonation affect the interaction.

Finally, we have also provided further insight into the structural basis for the previously reported high binding affinity of TDP-43 CTD to parallel G4 quadruplex structures (55–58), observing that contributions from aromatic residues to RNA binding dramatically increase in the presence of quadruplex structures. Interestingly, we also demonstrate that this unique binding interaction is not G4-specific, as the protein displayed a nearly identical binding properties with I-motif structures formed by two distinct sequences. Previous claims that the binding interaction between TDP-43 CTD strongly prefers to bind parallel G4 over antiparallel G4 structures (57) would be interesting to probe with these methods. Since we did not characterize the interaction of the CTD with an anti-parallel G4 structure during this study, we cannot rule out the possibility that TDP-43 CTD would display a different interaction with this structure, but this would then suggest that it is the previously observed lack of interaction with anti-parallel G4 structures that is unique, rather than the strong interactions with parallel G4 structures.

Overall, our findings suggest that the CTD contributes to the UG-rich binding preference reported for TDP-43 RRMs (34,35), potentially contributing to the specificity of TDP-43 for UG-rich RNAs previously ascribed to sequence preference only of the RRMs. Importantly, the primary binding region in TDP-43 CTD is directly adjacent to the RRMs, suggesting they may work together to recognize RNA partners, similar to the contacts made by the disordered and folded RNA binding domains of FUS (14). In the future we aim to investigate how the CTD and RRMs work together to recognize and bind the RNA targets of TDP-43, to provide a more holistic model of RNA recognition by TDP-43. Furthermore, given that many hnRNPs contain disordered regions enriched in aromatic and positively charged residues, we hypothesize that many hnRNPs may also display a binding preference for U-rich and G-rich sequences. More studies investigating the contributions of the disordered region of hnRNPs to RNA binding will be needed to better understand the generalizability of these findings.

## Acknowledgements

NMR experiments were conducted with the support of the Structural Biology Core Facility in the Division of Biology and Medicine at Brown University and with assistance from Mandar Naik. All-atom MD simulations were conducted with the advanced computing resources provided by Texas A&M High Performance Research Computing.

## Author Contributions

R.Z.P., H.L.D., V.J., and N.L.F. designed and analyzed the biochemical and biophysical experiments, and R.Z.P., H.L.D., and V.J. performed the experiments. B.O., Q.C., P.M., and J.M. designed, performed and analyzed data for MD simulations. R.Z.P., B.O., P.M., J.M. and N.L.F wrote the manuscript with contributions from all authors.

## Supplementary Data Statement

Supplementary Data are available at *NAR* Online

## Data Availability

NMR chemical shift assignments for WT TDP-43 and variants can be obtained online from the Biological Magnetic Resonance Database (Submission in progress) (BMRB, http://www.bmrb.wisc.edu/). Plasmids generated herein can be found at Addgene.org

