## Supplemental Figures for "The Structural Basis for RNA Binding and Recognition of the Disordered Prion-Like Domain of TDP-43"

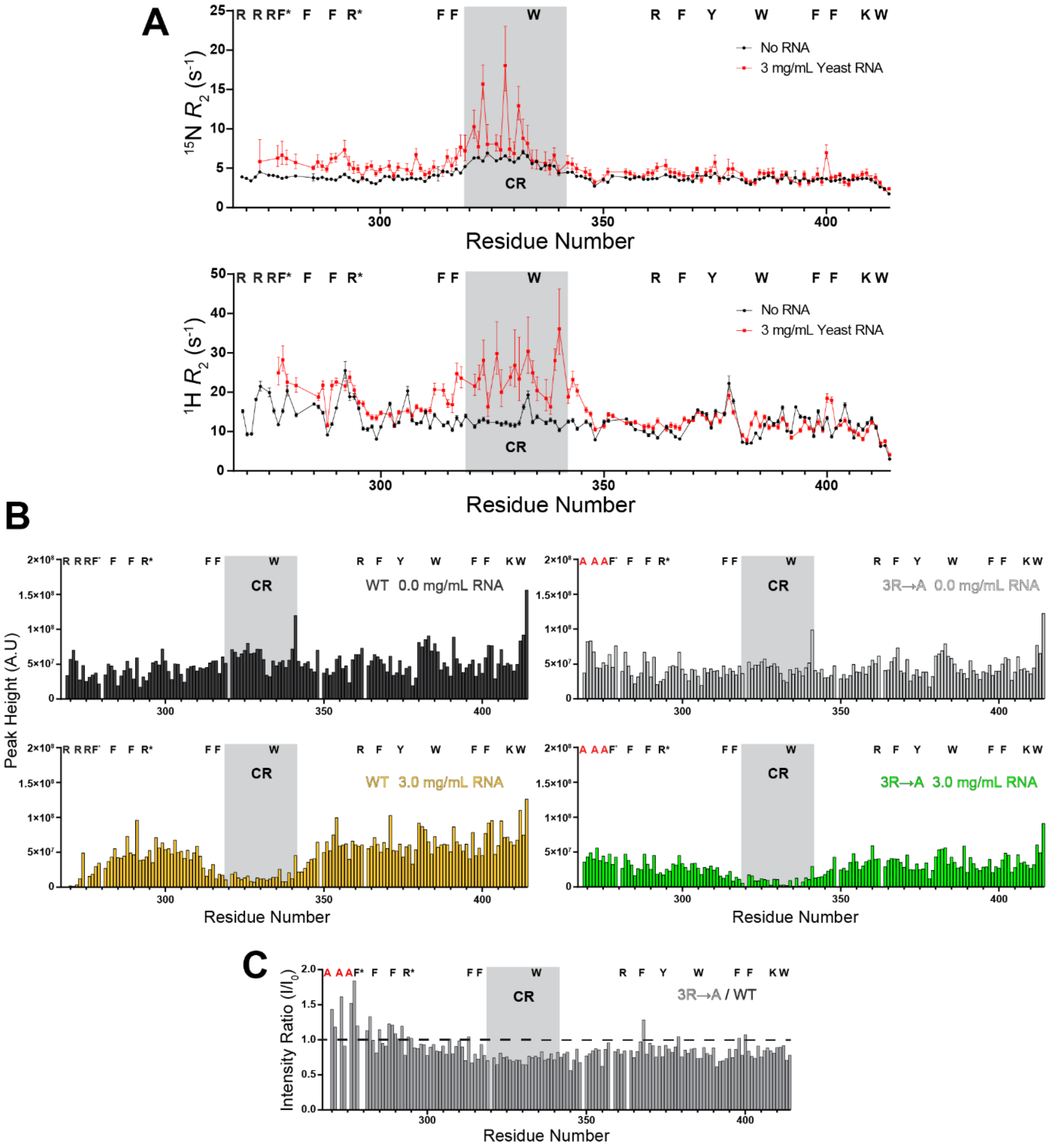

**Figure S1: RNA Interactions and N-terminal Arginine Mutations Change the Peak Intensity and Transverse Relaxation of the CTD: (A)** $^{15}$N and $^{1}$H $R_2$ ($s^{-1}$) analysis of TDP-43 CTD as a function of residue position in the absence (black) and presence of 3 mg/mL Torula Yeast RNA (red). $^{15}$N $R_2$ increased in the N-terminal binding region and the conserved region in the presence of RNA. $^{1}$H $R_2$ increased in the conserved region but sometimes decreased in the disordered regions. Additionally, $^{1}$H $R_2$ was observed to be higher in the Arg rich N-terminus than in other structural regions before RNA binding most likely due to chemical exchange events mediated by the N-terminal Arg. **(B)** Peak heights of WT TDP-43 CTD and 3R→A variant in the absence (black and grey respectively) and presence of 3 mg/mL yeast RNA (gold and green) displaying that for the WT residues that were initially low in intensity increased in intensity upon addition of RNA. **(C)** Peak intensity ratio of the 3R→A variant normalized to the WT displaying that the intensity of N-terminal residues increased in intensity upon the loss of Arg residues suggesting they mediate some form of  chemical exchange that gives rise to higher relaxation rates in the WT, which are then disrupted by their interactions with RNA.

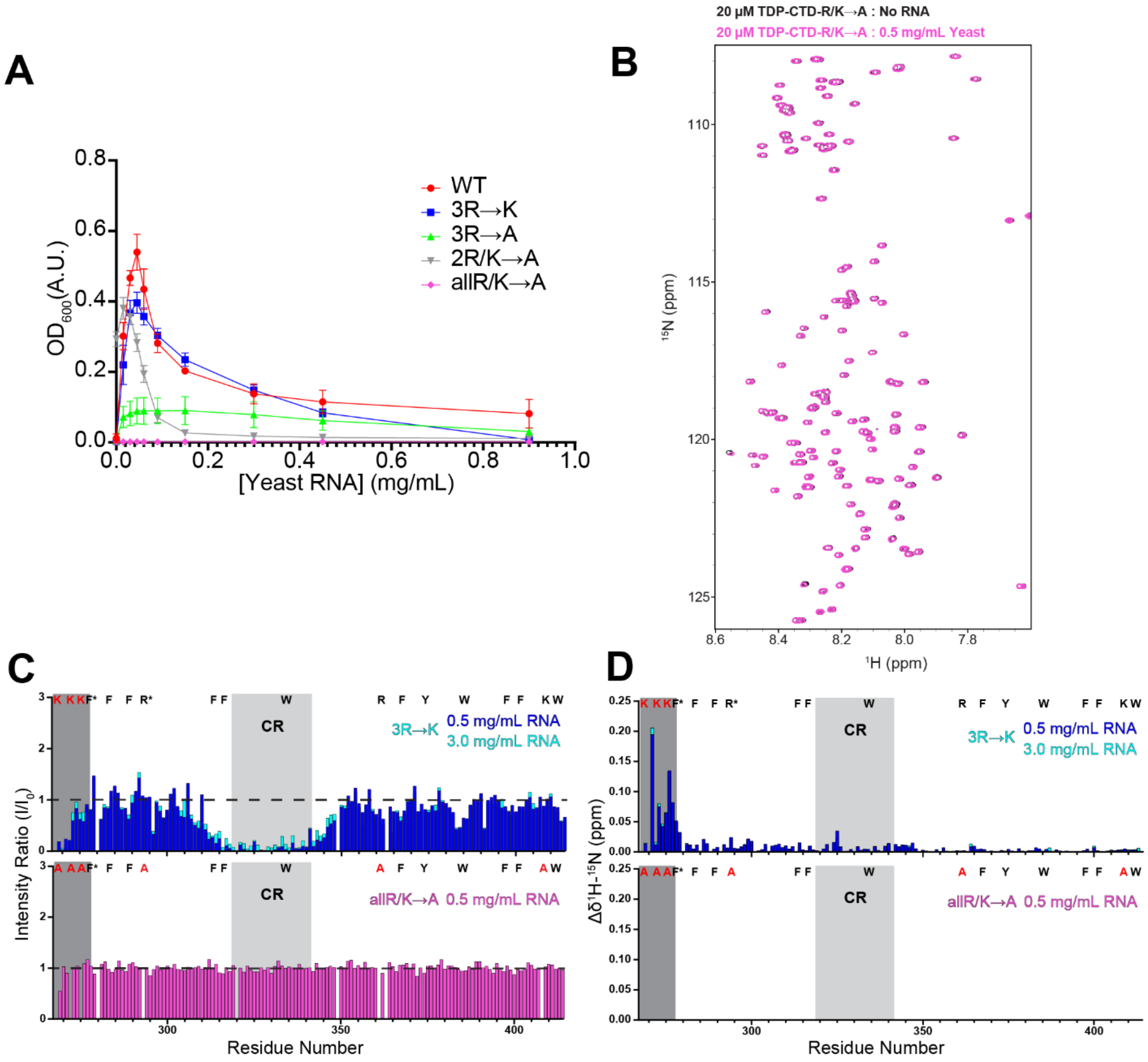

**Figure S2: Mutation of Arginine Residues Dramatically Affect RNA Induced LLPS and RNA Interaction: (A)** Turbidity assay comparing phase separation, measured as 600 nm optical density, of 20 μM WT TDP-43 CTD (red), TDP-43 CTD 3R→K (blue), TDP-43 CTD 3R→A (green), TDP-43 CTD 2R/K→A (grey), and TDP-43 CTD allR/K→A (purple), as a function of Torula Yeast RNA concentration (20 mM MES pH 6.1, 160 mM Urea) demonstrating how different charged residues impact RNA mediated phase separation. **(B)** $^{1}H$-$^{15}N$ HSQC overlay of 20 μM WT TDP43-CTD allR/K→A (20 mM MES pH 6.1) in the absence (black) and presence of 0.5 mg/mL Torula Yeast RNA (magenta) displaying no change in the protein spectra upon RNA addition. **(C)** Normalized NMR peak intensity ratios comparing the change in intensity of 20 μM TDP-43 CTD 3R→K titrated with 0.5 mg/mL Torula Yeast RNA and 3.0 mg/mL Torula Yeast RNA and TDP-43 CTD allR/K→A titrated with 0.5 mg/mL Torula Yeast RNA demonstrating how mutations change TDP-43 CTD RNA binding interactions. **(D)** Combined chemical shift perturbation plots of 20 μM TDP-43 CTD 3R→K titrated with 0.5 mg/mL Torula Yeast RNA and 3.0 mg/mL Torula Yeast RNA and TDP-43 CTD allR/K→A titrated with 0.5 mg/mL Torula Yeast RNA demonstrating how arginine mutations change the number and magnitude of RNA induced CSPs.

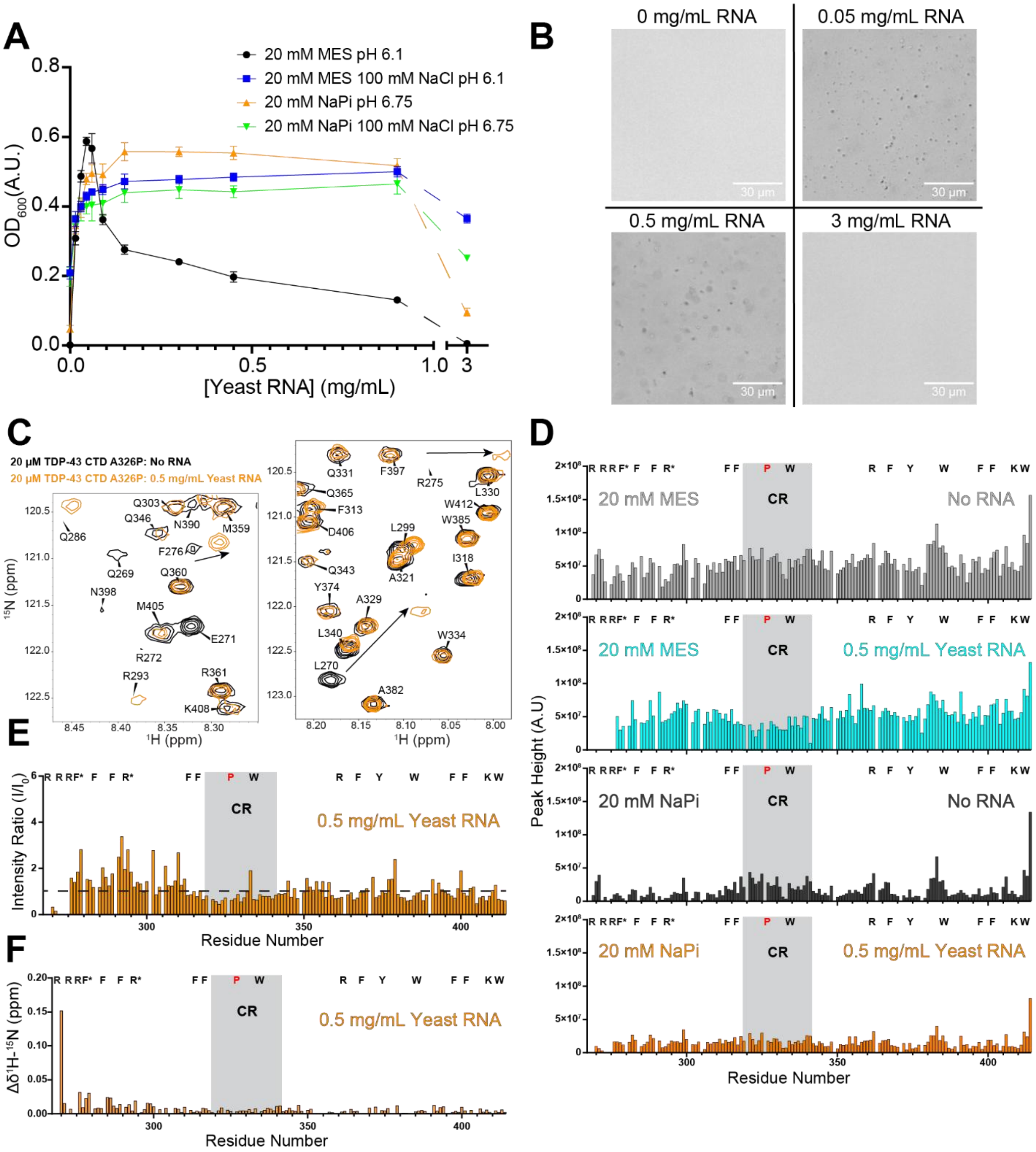

**Figure S3: Effects of Buffer on Reentrant LLPS and RNA Induced NMR Spectral Changes of TDP43 CTD: (A)** Turbidity assay comparing phase separation, measured as 600 nm optical density of 20 μM WT TDP-43 CTD as a function of Torula Yeast RNA concentration in several different buffer systems including 20 mM MES pH 6.1 (black), 20 mM MES 100 mM NaCl pH 6.1 (blue), 20 mM NaPi pH 6.75 (orange), and 20 mM NaPi 100 mM NaCl pH 6.75 (green). The presence of ionic species does not disrupt RNAs ability to induce phase separation but increase the amount of RNA required for reentrant behavior. **(B)** DIC micrographs of 20 µM WT TDP-43 CTD in the absence and presence of 0.05 mg/mL, 0.5 mg/mL, and 3 mg/mL of Torula Yeast RNA (20 mM NaPi pH 6.75, 160 mM Urea). **(C)** $^{1}$H-$^{15}$N HSQC overlay ROIs of 20 µM TDP-43-A326P in the absence (black) and presence of 0.5 mg/mL Torula Yeast RNA (orange) displaying the appearance of peaks in the presence of RNA and strong chemical shift perturbations. **(D)** Resonance intensities of 20 µM TDP-43-A326P CTD reported by NMRFAM Sparky in the absence (grey) and presence of 0.5 mg/mL Torula Yeast RNA (light blue) buffered by 20 mM MES pH 6.1 or in the absence (black) and presence of 0.5 mg/mL Torula Yeast RNA (orange) buffered by 20 mM NaPi pH 6.75. **(E)** Normalized NMR peak intensity ratios comparing the change in intensity of 20 µM TDP-43 CTD A326P residues (20 mM sodium phosphate buffer, pH 6.75) in the presence of 0.5 mg/mL Torula Yeast RNA (orange). N-terminal residues display an increase in intensity in the presence of RNA similar to spectra collected with 20 mM MES pH 6.1. **(F)** Combined $^{1}$H -$^{15}$N NMR chemical shift perturbations of 20 µM TDP-43 CTD A326P CTD residues (20 mM sodium phosphate buffer, pH 6.75) in the presence of 0.5 mg/mL Torula Yeast RNA (orange) displaying that CSPs still occur at N-terminal region.

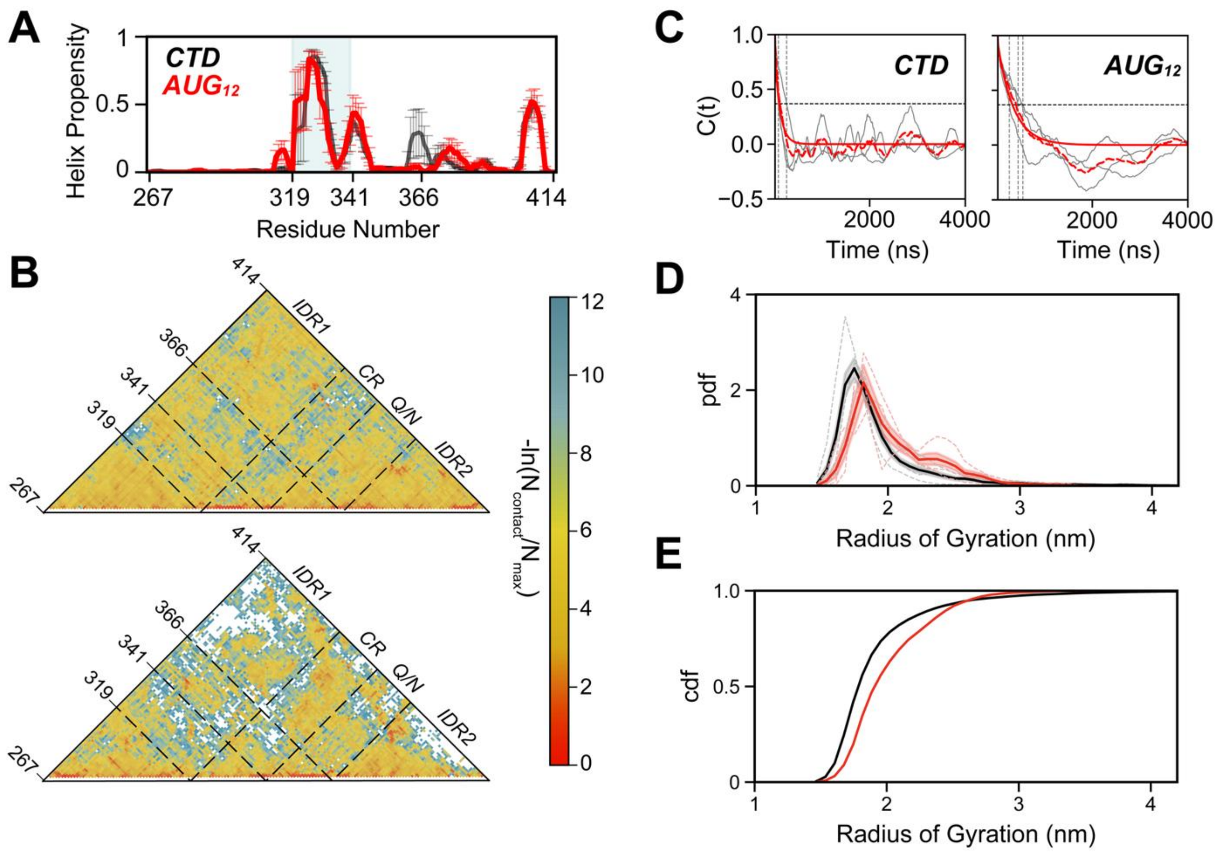

**Figure S4: Effect of AUG12 Binding on the CTD Conformational Ensemble. (A)** α-helix propensity across the CTD sequence. CR region indicated by gray shading. Data represents mean across three independent simulations with error bars representing standard error of the mean. **(B)** Intramolecular contact map showing heavy atom pairs within 4.5 Å, calculated after excluding the first 500 ns and normalized by frame number. Upper diagonal: without RNA; lower diagonal: with AUG12. RNA binding disrupts contacts between IDR1 (N267–I318) and IDR2 (A366–M414). **(C)** Radius of gyration autocorrelation function C(t) for CTD with and without RNA. Data show three independent replicas. Vertical dotted lines indicate where C(t) crosses 1/e (≈ 0.368). Mean crossing times: 128 ns (without RNA), 406 ns (with RNA). Based on these values, 500 ns was selected as equilibration time for all systems. **(D)** Radius of gyration probability density (pdf) distributions for CTD without RNA (black) and with AUG12 (red). Individual replicas (dashed lines) and mean distribution (solid lines) are shown. **(E)** Cumulative probability (cdf) of mean CTD Rg with (red) and without (black) $AUG_{12}$.

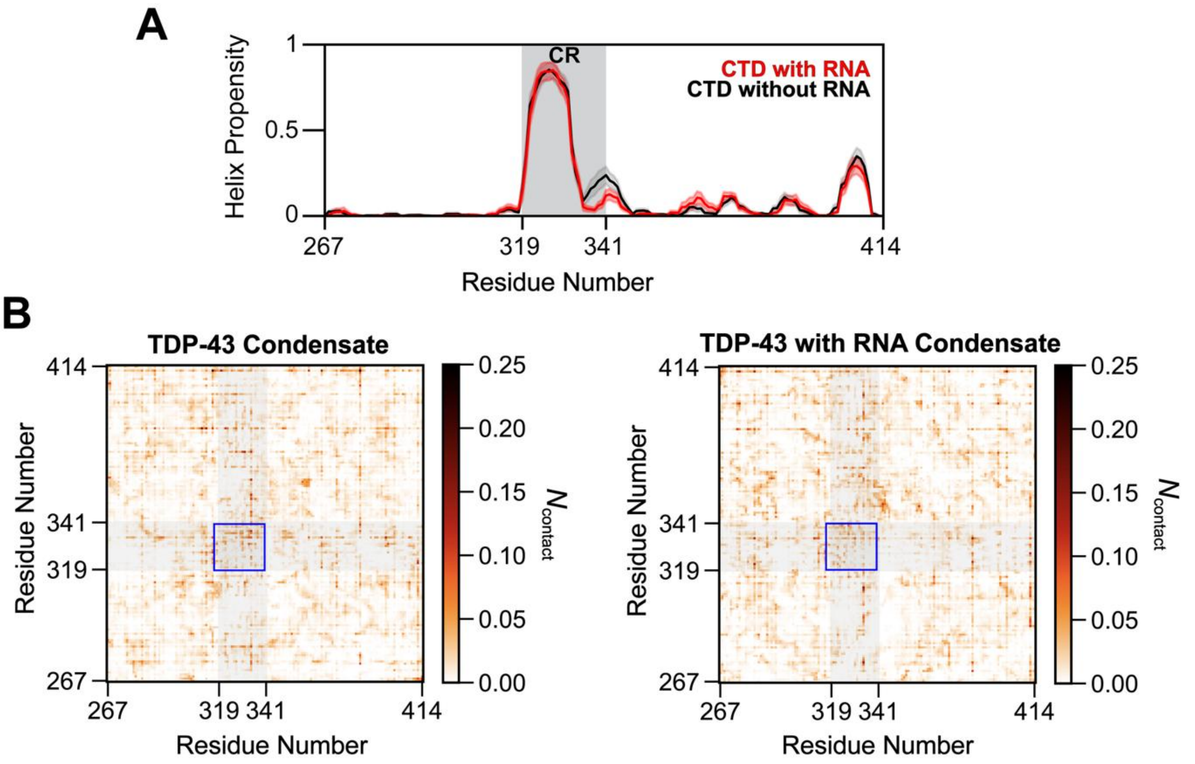

**Figure S5: Conserved Region Helicity and Interactions Remain Similar within the Dense Phase Regardless of RNA (A)** α-helix propensity across the CTD sequence in the dense phase without RNA (black) and with RNA (red). CR region indicated by gray shading. Data represent mean across protein chains; error bars indicate standard error of the mean. **(B)** Intermolecular contact maps between CTD chains, calculated as heavy atom pairs within 4.5 Å and normalized by the number of possible chain pairs (40 × 39 / 2) and frames. Top: without RNA; bottom: with RNA. Blue box indicates CR-CR interactions between different CTD chains.

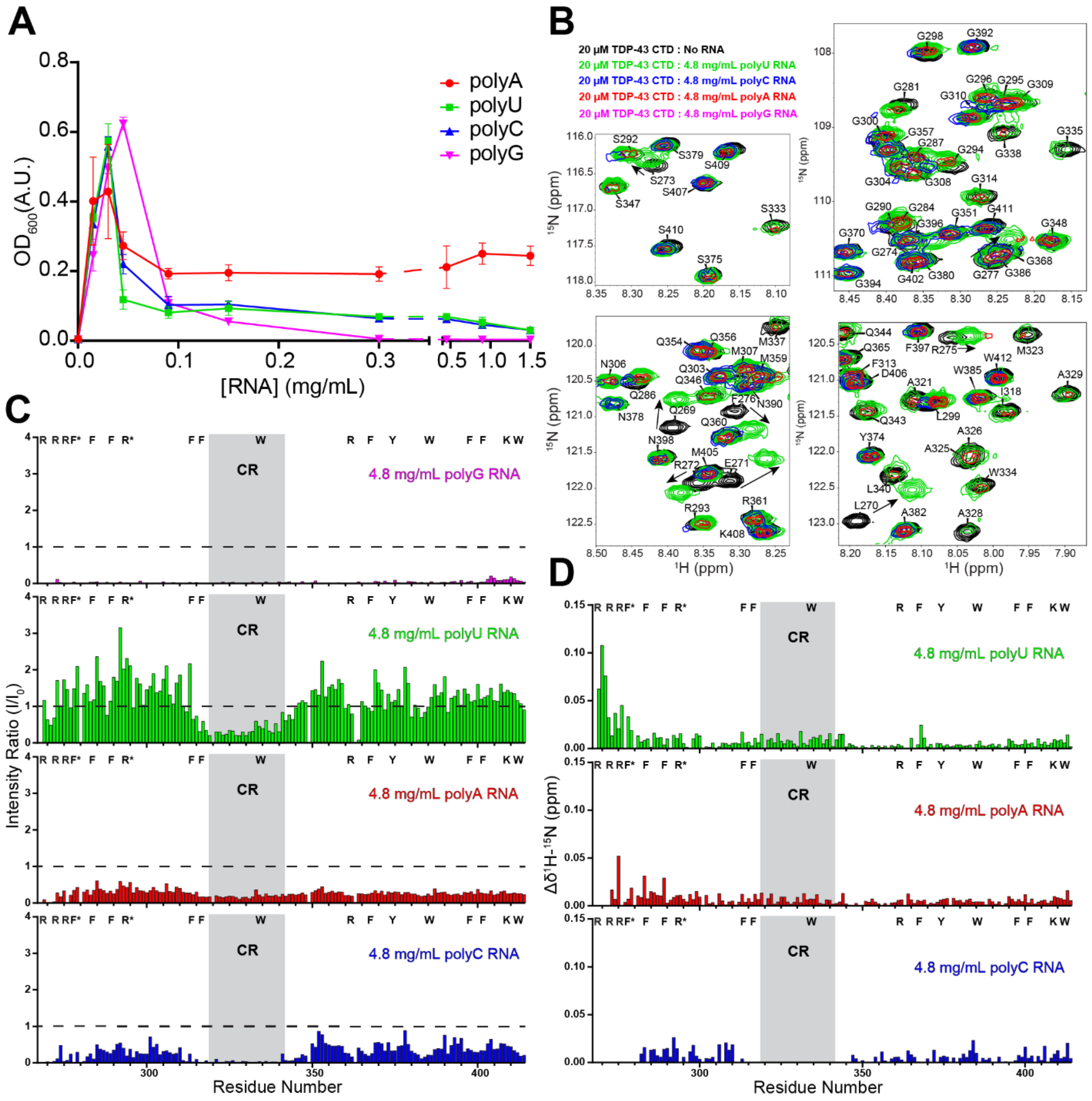

**Figures S6: TDP-43 CTD Displays Differences in Phase Separation and RNA Binding Interaction Depending on Nucleotide Base: (A)** Turbidity assay comparing phase separation, measured as 600 nm optical density of 20 µM WT TDP-43 CTD as a function of polyA, polyU, polyC, and polyG RNA concentration (20 mM MES pH 6.1) displaying differences in reentrant behavior depending on nucleotide base present in homopolymer. **(B)** $^{1}$H-$^{15}$N HSQC overlay ROIs of 20 µM WT TDP-43 CTD in the absence (black) and presence of 4.8 mg/mL polyU RNA (green), 4.8 mg/mL polyC RNA (blue), 4.8 mg/mL polyA RNA (red), and 4.8 mg/mL polyG RNA (magenta). **(C)** Normalized NMR peak intensity ratios comparing the change in intensity of 20 µM WT TDP-43 CTD residues (20 mM MES, pH 6.1) in the presence of 4.8 mg/mL polyG RNA (magenta), 4.8

mg/mL polyU RNA (green), 4.8 mg/mL polyC RNA (blue), and 4.8 mg/mL polyA RNA (red) displaying distinctly different intensities depending on homopolymer. **(D)** Combined $^{1}H$-$^{15}N$ NMR chemical shift perturbations of 20 μM WT TDP-43 CTD residues (20 mM MES, pH 6.1) in the presence of 4.8 mg/mL polyU RNA (green), 4.8 mg/mL polyA RNA (red), and 4.8 mg/mL polyC RNA (blue) displaying large CSPs in the presence of polyU compared to polyC and polyA.

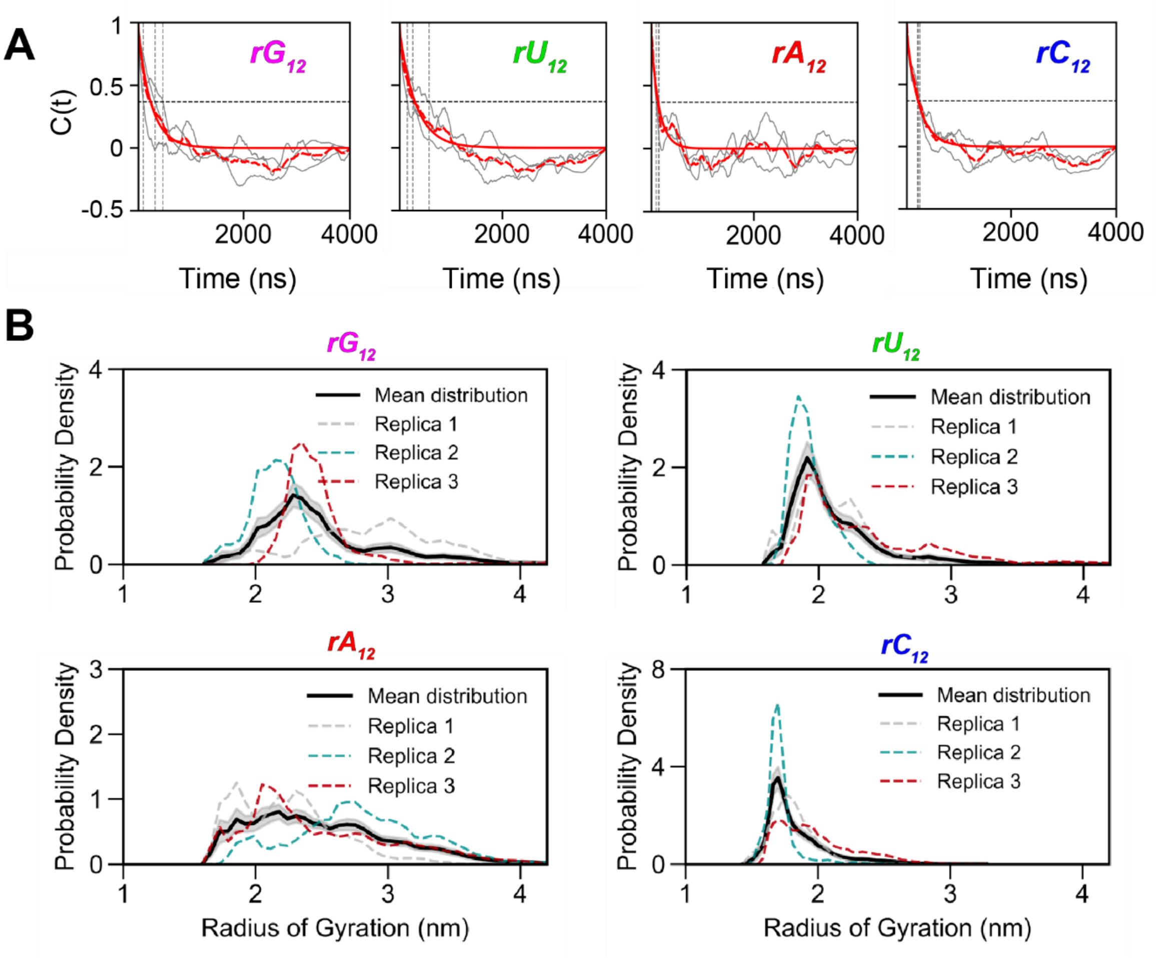

**Figure S7: Conformational Sampling of CTD with Homopolymeric RNA Sequences.** **(A)** Radius of gyration autocorrelation function C(t) for CTD in the presence of $rG_{12}$ (magenta), $rU_{12}$ (green), $rA_{12}$ (red), and $rC_{12}$ (blue). Data show three independent replicas for each sequence. Vertical dotted lines indicate where C(t) crosses 1/e (≈ 0.368). Based on these crossing times, 500 ns was selected as equilibration time for all systems. **(B)** Radius of gyration distributions for each RNA sequence. Individual replicas (dashed lines) and mean distribution (solid black) are shown.

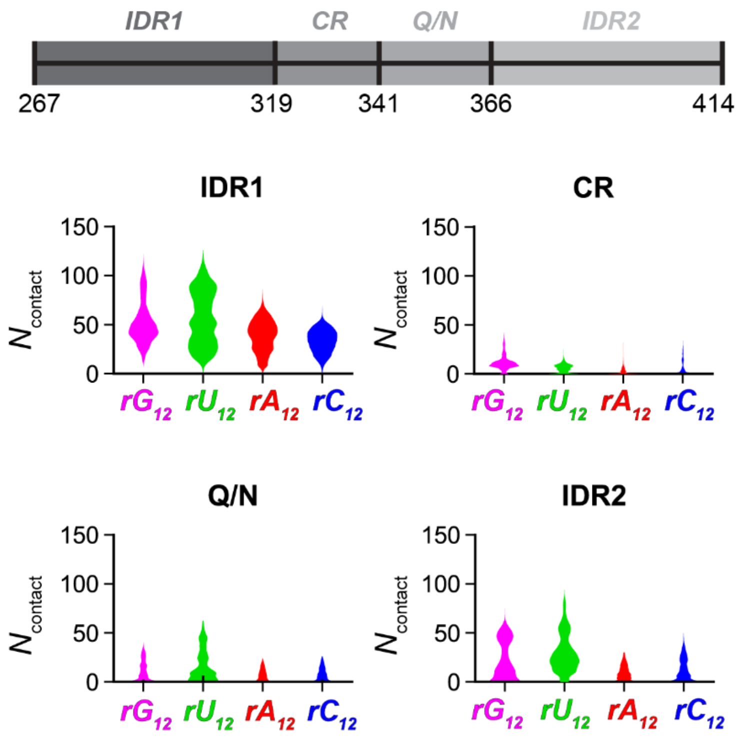

**Figure S8: CTD-RNA Interactions in the Presence of Homopolymeric Sequences.** Top: Schematic of different regions within CTD. Bottom: Distribution of protein-RNA contacts by region (IDR1: 267–318, CR: 319–341, Q/N: 342–365, IDR2: 366–414) for each homo-oligomer. Violin plots show contact distributions across trajectory frames.

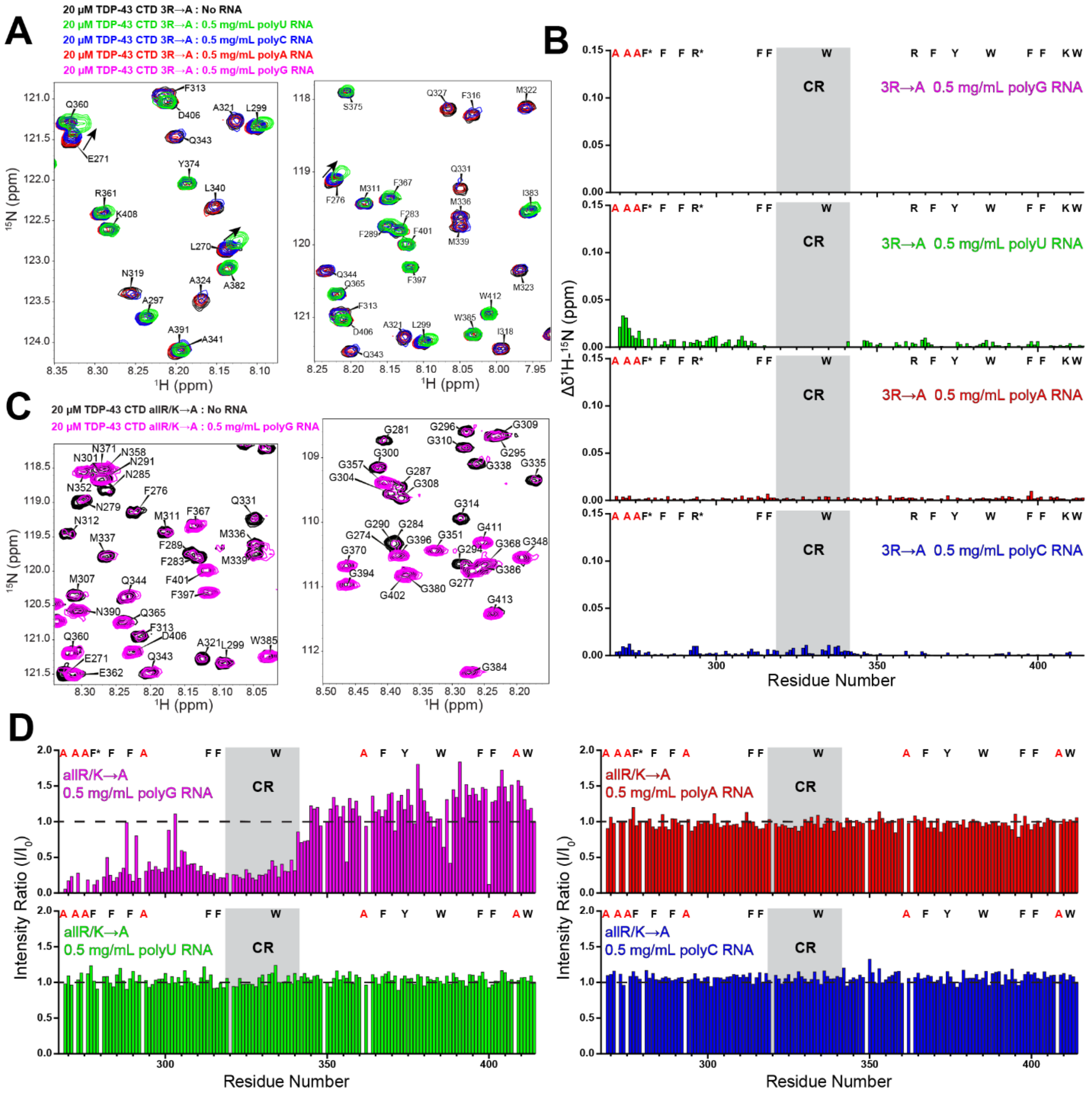

**Figure S9: Alanine Screening Displays Difference in Homopolymer Interactions Depending on Removal of Charged Residues: (A)** $^{1}$H-$^{15}$N HSQC overlay ROIs of 20 µM TDP-43 CTD 3RtoA (20 mM MES pH 6.1) in the absence (black) and presence of 0.5 mg/mL polyU (green), polyC (blue), polyA RNA (red), and polyG RNA (magenta) displaying differences in peak intensity for all RNAs and weak CSPs with polyU. **(B)** Combined $^{1}$H -$^{15}$N NMR chemical shift perturbations of 20 µM TDP-43 CTD 3R→A residues (20 mM MES, pH 6.1) in the presence of 0.5 mg/mL polyG RNA (magenta), polyU (green), polyC (blue), and polyA RNA (red) displaying the largest CSPs occurred in the presence of polyU RNA. **(C)** $^{1}$H-$^{15}$N HSQC overlay ROIs of 20 µM TDP-43 CTD allR/K→A (20 mM MES pH 6.1) in the absence (black) and presence of 0.5 mg/mL polyG

RNA (magenta) displaying signal attenuation for some residues. **(D)** Normalized peak intensity ratio plots of TDP-43 CTD allR/K→A in the presence of 0.5 mg/mL polyG RNA (magenta) polyU (green), polyA (red), and polyC RNA (blue) displaying reduction in peak intensity for IDR1 and conserved region in the presence of polyG RNA but no change for the other RNA homopolymers.

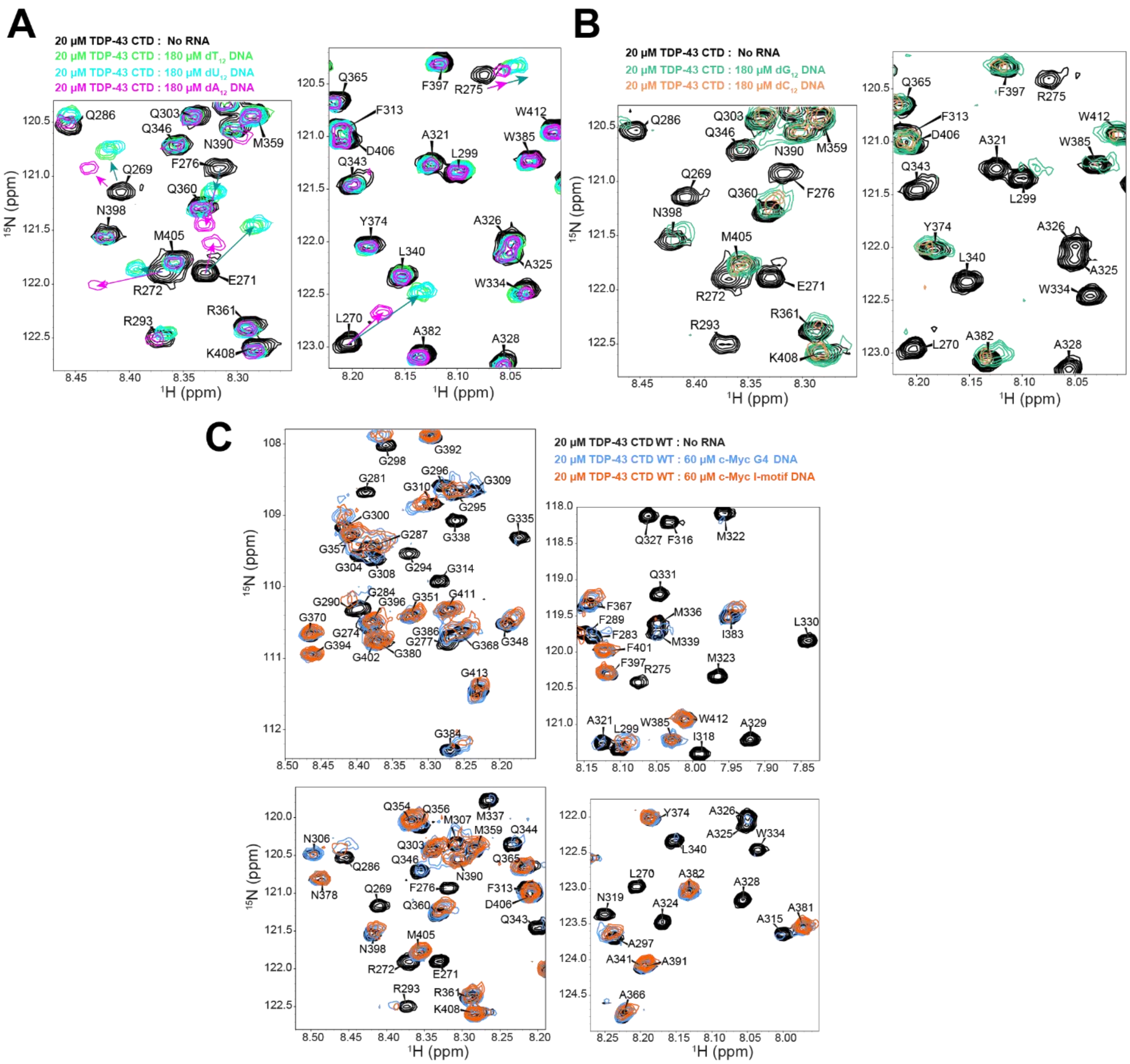

**Figure S10: TDP-43 CTD Displays Unique Interaction in the Presence of Quadruplex Structures:** **(A)** $^{1}$H-$^{15}$N HSQC overlay ROIs of 20 µM WT TDP-43 CTD in the absence (black) and presence of 180 µM $dT_{12}$ (light green), 180 µM $dU_{12}$ (light blue), and 180 µM $dA_{12}$ (magenta). TDP-43 CTD displays large CSPs in presence of each 12mer ssDNA but magnitude of CSPs differ depending on base. **(B)** $^{1}$H-$^{15}$N HSQC overlay ROIs of 20 µM WT TDP-43 CTD in the absence (black) and presence of 180 µM $dG_{12}$ (ocean green) and 180 µM $dC_{12}$ (orange). TDP-43 CTD displays loss of peaks in the presence of $dG_{12}$ and $dC_{12}$ 12mers. **(C)** $^{1}$H-$^{15}$N HSQC overlay ROIs of 20 µM WT TDP43-CTD (20 mM MES pH 6.1) in the absence (black) and presence of 60 µM c-Myc G4 quadruplex DNA (sky blue), and 60 µM c-Myc I-motif DNA (orange). Loss of the same peaks with both ssDNA sequences demonstrates the protein interaction is similar for both quadruplex structures.

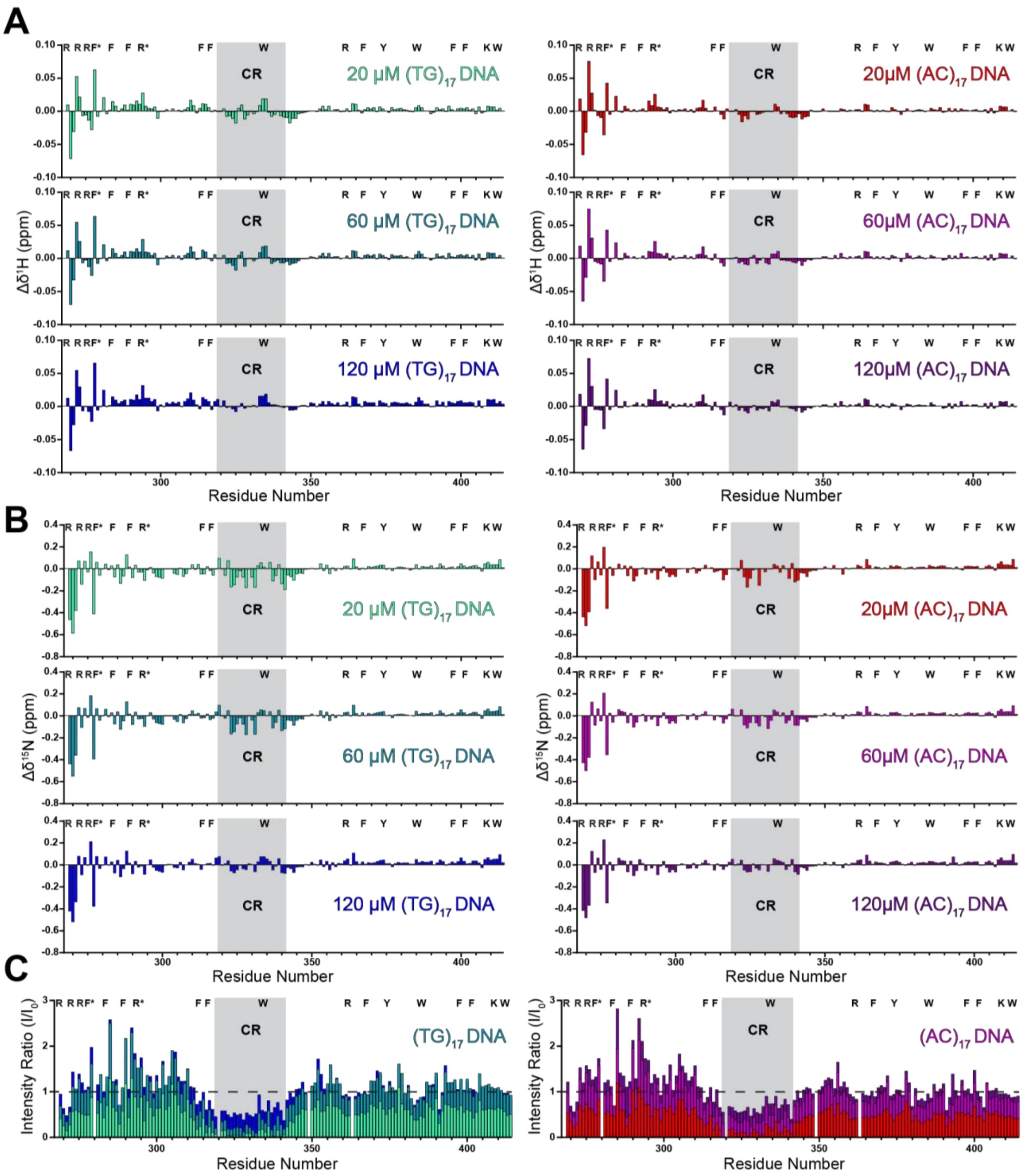

**Figure S11: ¹H and ¹⁵N Chemical Shift Perturbations and Intensity Changes of TDP-43 CTD Titrated with TG-Rich and AC-Rich Sequences: (A)** ¹H NMR chemical shift perturbations (CSPs) of 20 μM WT TDP-43 CTD (20 mM MES, pH 6.1) titrated with 20 μM (red), 60 μM (magenta), and 120 μM (purple) $(AC)_{17}$ DNA or 20 μM (aquamarine), 60 μM (light blue), and 120 μM (dark blue) $(TG)_{17}$ DNA. **(B)** ¹⁵N NMR chemical shift perturbations (CSPs) of 20 μM WT TDP-43 CTD (20 mM MES, pH 6.1) titrated with 20 μM (red), 60 μM (magenta), and 120 μM (purple) $(AC)_{17}$ DNA or 20 μM (aquamarine), 60 μM (light blue), and 120 μM (dark blue) $(TG)_{17}$ DNA. **(C)** Overlayed normalized NMR peak intensity ratios as a function of sequences position comparing the change in intensity of 20 μM WT TDP-43 CTD (20 mM MES, pH 6.1) titrated with 20 μM (red), 60 μM (magenta), and 120 μM (purple) $(AC)_{17}$ DNA or 20 μM (aquamarine), 60 μM (light blue), and 120 μM (dark blue) $(TG)_{17}$ DNA displaying differences in conserved region residue peak intensity depending on DNA base composition.
